# Post-transcriptional regulation of Nrf2-mRNA by the mRNA-binding proteins HuR and AUF1

**DOI:** 10.1101/565937

**Authors:** Jesse R. Poganik, Marcus J. C. Long, Yimon Aye

## Abstract

The Nrf2 signaling axis is a target of covalent drugs and bioactive native electrophiles. However, much of our understanding of Nrf2 regulation has been focused at the protein-level. Here we report a post-transcriptional modality to directly regulate Nrf2-mRNA. Our initial studies focused on the effects of the key mRNA-binding protein (mRBP) HuR on global transcriptomic changes incurred upon oxidant or electrophile stimulation. These RNA-sequencing data and subsequent mechanistic analyses led us to discover a novel role of HuR in regulating Nrf2 activity, and in the process, we further identified the related mRBP AUF1 as an additional novel Nrf2 regulator. Both mRBPs regulate Nrf2 activity by direct interaction with the Nrf2 transcript. Our data showed that HuR enhances Nrf2-mRNA maturation and promotes its nuclear export; whereas AUF1 stabilizes Nrf2-mRNA. Both mRBPs target the 3ʹ–UTR of Nrf2-mRNA. Using a Nrf2-activity reporter zebrafish strain, we document that this post-transcriptional control of Nrf2 activity is conserved at the whole-vertebrate level.

## INTRODUCTION

The Nrf2 signaling axis is the cell’s linchpin stress defense^1^, whereby the central player, the Nrf2 transcription factor, controls the expression of numerous genes for cytoprotection and detoxification^2^. Nrf2 upregulation accompanies cell stimulation with reactive electrophilic and oxidative species (RES/ROS), which are increasingly appreciated as bona fide cell signaling cues^3^. Upon stimulation with either endogenous or environmental RES/ROS, the cell mounts a stress defense through upregulation of Nrf2 pathway. Beyond RES/ROS-stimulated conditions, Nrf2 activity at basal (i.e., non-stimulated) conditions is tightly regulated. However, the most well-understood modes of Nrf2 regulation—both at basal and stimulated cell states—are centered around Nrf2-protein-level control^2^. Among the most notable players are Keap1 and β-TrCP, which independently maintain low steady-state levels of Nrf2-protein in non-stimulated cells through Nrf2-proteasomal targeting^4,5^. Deregulation of protein-level control of Nrf2 transcriptional activity is common in various cancers^6^. Recent efforts to modulate Nrf2 signaling by targeting protein-based Nrf2-regulators like Keap1 have underscored the pharmaceutical relevance of this pathway^7^. In addition to protein-level regulation of Nrf2, some microRNAs (miRNAs) target this pathway either by direct Nrf2-targeting, or targeting of other protein-regulators such as Keap1^8^. However, expression of these miRNAs tends to be highly tissue/context-specific. A *general* regulatory mechanism (e.g., through interaction with a ubiquitously-expressed protein) of Nrf2 activity at the mRNA level has not been reported.

Regulation of mRNA is a complex process that can be mediated through structural^9^, sequence^10^, or epitranscriptomic^11^ elements within a transcript. The ubiquitously-expressed mRNA-binding protein (mRBP) HuR (ELAVL1) is a postulated stress-relevant protein that binds to AU-rich sites within regulatory mRNA targets and regulates their expression via post-transcriptional mechanisms^12^. These binding sites typically reside within 3ʹ–untranslated regions (UTRs) of target transcripts. However, binding within introns, coding regions, and 5ʹ–UTRs has also been observed. Regulation of mRNA targets by HuR can occur via direct binding, or indirectly by miRNA-dependent mechanisms^13,14^.

State-of-the-art sequencing techniques such as PAR-CLIP^15,16^ have revealed thousands of functionally-diverse targets of HuR^17,18^. Although Nrf2-mRNA has been detected by PAR-CLIP analysis of HuR, no functional validations nor interaction studies have been made, likely because the Nrf2 transcript appears as a low-frequency hit [with only 46 T-to-C/A-to-G conversions for HuR (marking crosslinks to HuR), compared to hundreds to thousands of conversions for the most highly-ranked transcripts such as AKT3 and POLA1]. However, PAR-CLIP rankings generally correlate poorly with HuR target affinity (Table S1); PAR-CLIP conversion number may be affected by varying expression levels of different targets or artifacts of the PAR-CLIP procedure. Accordingly, there remains a need to mechanistically investigate how HuR regulates important disease-relevant targets such as Nrf2-mRNA.

In addition to the difficulty in ranking importance of hits from high-throughput data sets, the multimodal regulatory activities of HuR render predicting functional consequences of a HuR-mRNA binding event difficult. HuR—distributed between the nucleus and cytosol in 10:1 ratio in HeLa cells^19^— largely modulates its target transcripts through alterations in mRNA-stability, typically by stabilizing bound transcripts^20^. Additional regulatory mechanisms employed by HuR on its target transcripts have also been reported: e.g., nuclear export of the cyclooxygenase COX-2 mRNA^21,22^ and splicing regulation of the death receptor FAS and the translational regulator eIF4Enif1^23,24^. However, the generality of these nuanced regulatory roles remains largely unclear, and their importance in Nrf2 regulation is completely unknown.

HuR is strongly linked to disease as a key player in inflammation and cancers among other disorders^25^. The growing appreciation of HuR as a major player in disease is underscored by recent efforts to screen for inhibitors of the HuR–RNA interaction^26–28^. The non-covalent inhibitors identified from such screens disrupt target-transcript binding or HuR oligomerization (key to HuR function). Some HuR inhibitors have shown promise in arresting cell cycle and inducing apoptosis in cultured lung cancer cells^29^. The mRNA-stabilizing ability of HuR is also affected by small-molecule stress-inducers such as the electrophilic prostaglandin A2 (PGA2) and lipopolysaccharide (LPS), which triggers oxidative stress^30–33^. However, no study has directly compared the role of HuR regulation of the Nrf2 signaling axis under conditions of oxidative or electrophilic stress.

Understanding HuR-regulatory circuits is further complicated by the presence of multiple mRBPs that bind to target mRNAs at the same/similar sites as HuR, but generally elicit antagonistic outputs to HuR binding. One such mRBP is AUF1 (HNRNPD), which shares significant target overlap with HuR based on PAR-CLIP analysis^34^. Mechanistically, in contrast to the canonical role of HuR in promoting mRNA stability, AUF1 typically promotes degradation of target mRNAs^35^. However, other regulatory mechanisms employed by AUF1 have recently come into focus: AUF1 can promote translation of myocyte enhancer factor MEF2C, but suppresses translation of profilin 1^34,36^. Crosstalk between HuR and AUF1 in regulating common mRNA targets has also been investigated for the cyclin-dependent kinase inhibitor p21 and cyclin D1^19^. Stability of both of these targets was regulated positively by HuR and negatively by AUF1. However, general understanding of these co-regulatory events by these two mRBPs remains limited. Whether HuR and AUF1 bind to different sites or compete for the same sites within a specific co-regulated transcript also remains unclear^19^. Nevertheless, AUF1 generally functions as the antipode of HuR in terms of target transcript regulation.

Herein, we describe novel post-transcriptional regulatory modes of the Nrf2 pathway, controlled by HuR and AUF1. Our initial RNA-sequencing analysis of HuR-dependent global transcriptomic changes in response to cell stimulation by H_2_O_2_ and 4-hydroxynonenal (HNE)—the prototypical oxidant (ROS) and electrophile (RES)—revealed that only the electrophile HNE causes significant induction of Nrf2-driven genes. Surprisingly, we found that HuR depletion generally diminished Nrf2 transcriptional activity in non-stimulated cells. Yet, upon electrophile stimulation, Nrf2 activity was more strongly upregulated in HuR-depleted cells. Our RNA-seq data implicate the involvement of different subsets of Nrf2-driven genes underlying these two effects. Our further investigations into Nrf2 regulation by HuR in non-stimulated cells led us to discover a previously-unrecognized regulatory loop controlling Nrf2 activity, which our data show is conserved from cultured cells to zebrafish. This newly-identified post-transcriptional regulatory mode of Nrf2 activity potentially offers a novel alternative intervention to modulate Nrf2/AR in disease.

## RESULTS

### HuR depletion perturbs the global transcriptomic status in electrophile-stimulated and non-stimulated cells

HuR has been implicated as a stress-responsive protein: cytoplasmic translocation of HuR is promoted by several small-molecule stressors, e.g., H_2_O_2_, arsenite, and the cyclopentenone prostaglandin A2 (PGA2), a Michael acceptor RES^30,31^. Treatment with PGA2 also increases the affinity of HuR for p21 mRNA^30^. Intrigued by reports of the stress-relevance of HuR, we launched gene-expression profiling studies in HuR-knockdown cells, in the presence or absence of small-molecule stress signals. HNE—a native RES with a reactive core chemically similar to that of PGA2—was selected as a representative bioactive RES^37,38^ and H_2_O_2_ was selected as a representative ROS. HuR-knockdown HEK293T cells were first generated (Table S2). Relative to a non-targeted shControl, HuR was knocked down by 70% in these cells (Figure S1A). We then treated these shControl and shHuR cells with HNE (25 μM, 18 h) or H_2_O_2_ (225 μM, 18 h) and subsequently sequenced their RNA, post ribosomal RNA (rRNA) depletion. The chosen concentrations of HNE/H_2_O_2_ correspond to the approximate EC_75_ for growth inhibition, measured over 48 h (Figure S2). Significant differential expression was evaluated with CuffDiff^39^, wherein gene-level pairwise comparisons having q-value (p-value corrected for multiple tests) < 0.05 are considered significantly differentially expressed (SDE).

Compared to non-treated control, HNE-stimulated shControl cells showed considerable upregulation of 11 genes out of 14 total genes SDE (79%). The data skewed in the positive direction overall, indicating a general stimulation of gene transcription [Figures 1A (data in blue) and S3; Tables S3 and S4]. In contrast, HNE-treated shHuR cells (compared to non-stimulated shHuR cells) showed upregulation of only 58% of the 12 genes SDE, and the dataset skewed in the negative direction overall [Figures 1A (data in red) and S3; Tables S3 and S4]. Thus, depletion of HuR compromises the ability of cells to mount positive transcriptional responses following electrophile stimulation. By contrast, neither shHuR nor shControl cells mounted as robust a response to H_2_O_2_ as to HNE. Compared to the number of genes SDE in HNE-stimulated shControl and shHuR cells (14 and 12, respectively), only 2 genes were SDE in H_2_O_2_-treated shControl cells, and no genes were SDE in H_2_O_2_-treated shHuR cells (Figure 1B; Table S3). Both control and HuR-knockdown datasets with H_2_O_2_-treatment skewed slightly in the negative direction (Figure S3; Tables S3 and S4).

**Figure 1.**
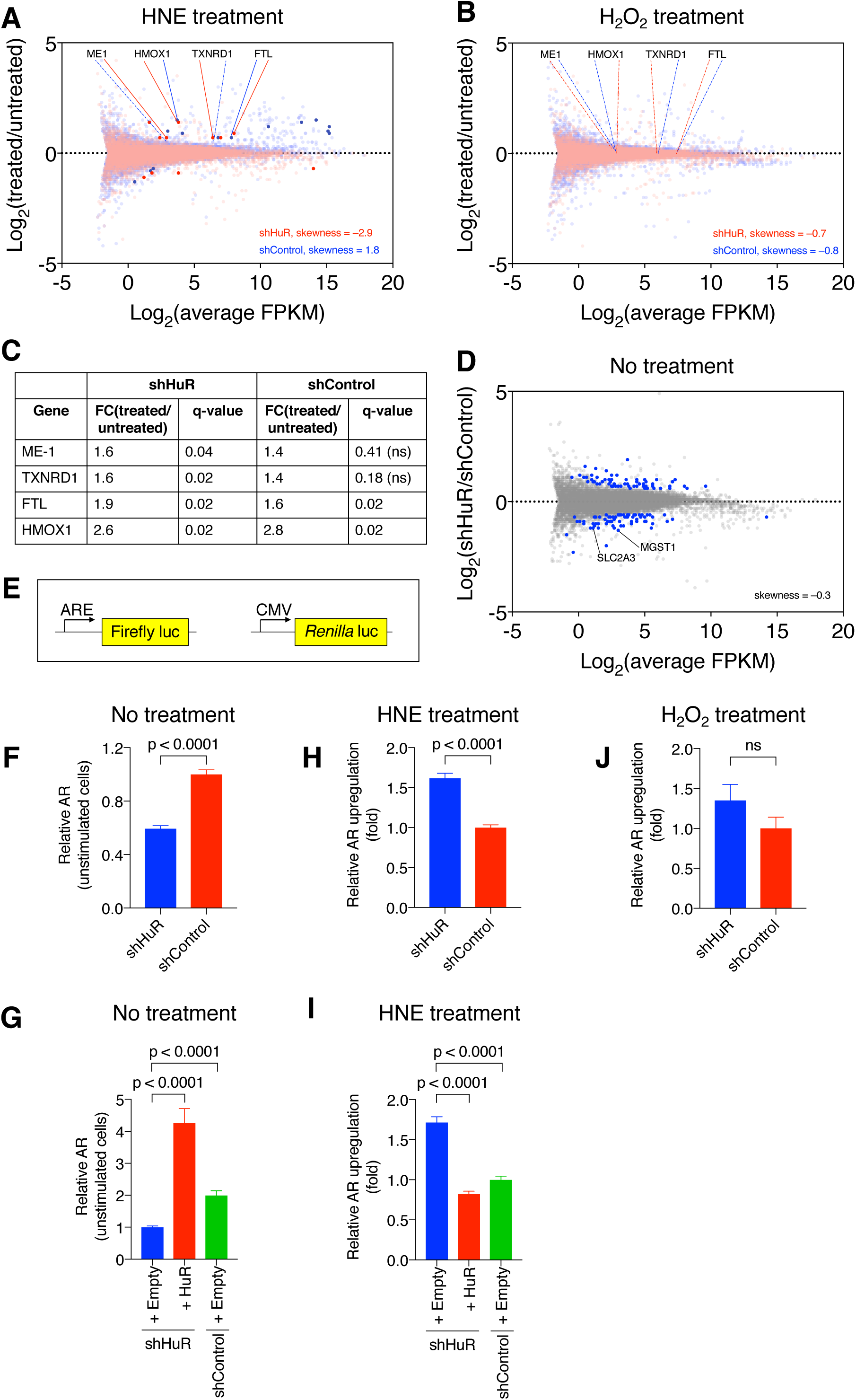
RNA-seq expression profiling indicates context-specific HuR-regulation in global and Nrf2-specific transcriptional activities. (**A** and **B**) Differential expression from RNA-seq analyses in shHuR or shControl HEK293T cells following HNE (25 uM, 18 h) or H_2_O_2_ (225 uM, 18 h) stimulation relative to respective non-stimulated cells. Genes significantly differentially expressed (SDE) are denoted with dark, opaque points; Nrf2-driven genes SDE in at least one comparison are labeled. Dashed lines to gene names indicate that the gene was not SDE in that comparison. See also Figures S1A, S2, S3, and Tables S2 and S3. (**C**) Fold changes (FCs) for selected Nrf2-driven genes of interest upon HNE stimulation relative to non-treated. Q-values were calculated with CuffDiff. See also Tables S3–S5. (**D**) Differential expression from RNA-seq analysis in non-stimulated shHuR cells relative to shControl cells. Genes SDE are denoted with blue points. Two Nrf2-driven genes found to be SDE are labeled. See also Tables S3–S5. (**E**) Nrf2 activity reporter system in which firefly luciferase is driven by Nrf2 and a constitutive *Renilla* luciferase is co-expressed as an internal normalization control. (**F**) Nrf2 activity of non-HNE-stimulated shHuR and shControl cells (mean±SEM, n=24). (**G**) Nrf2 activity was measured as in **F**, but ectopic HuR was expressed in non-HNE-stimulated shHuR cells (mean±SEM, n=7 for shHuR+HuR and n=8 for other conditions). See also Figure S4. (**H**) Nrf2 activity in HNE (25 μM, 18 h)-stimulated shHuR and shControl (mean±SEM, n=16)]. (**I**) Nrf2 activity was measured as in **H**, but ectopic HuR was expressed in HNE(25 μM, 18 h)-stimulated shHuR cells (mean±SEM, n=8). See also Figure S4. (**J**) Nrf2 activity in H_2_O_2_(225 μM, 18 h)-stimulated shHuR and shControl cells (mean±SEM, n=7 for shHuR and 8 for shControl)]. All p-values were calculated with Student’s T-test. For **A** and **B**, data were derived from n=2 independent biological replicates per treatment condition. Skewness was calculated with Prism.

### HuR is a context-specific regulator of Nrf2 transcriptional activity

Beyond HuR-dependent effects on global transcriptional status, we compared the expression of Nrf2-driven genes in shHuR and shControl cells following HNE stimulation. We identified one gene, PIR, that was not significantly upregulated in shHuR knockdown lines upon HNE treatment, but was upregulated in shControl lines upon HNE stimulation. However, we also detected a further 4 Nrf2-driven genes (ME-1, TXNRD1, FTL, and HMOX1) that were upregulated in both shHuR and shControl cells treated with HNE (Figures 1A and C). Intriguingly, 3 of these genes were upregulated to a greater extent in shHuR cells than in shControl cells: ME-1, TXNRD1, and FTL, showed 1.05–1.2-fold higher fold-change (FC)_(treated/untreated)_ in shHuR cells versus shControl cells (Figures 1A and C). This trend contrasts with the overall downregulation of global transcriptional activity following HNE-stimulation in shHuR-(0.98 average fold change, –2.9 skewness) compared to shControl-lines [1.0 average fold change, 1.8 skewness (Figures 1A and S3)]. We did not observe significant differential expression of these genes between *non-stimulated* shHuR and shControl cells (Figure 1D and Table S3), but we were able to identify three other Nrf2-driven genes (SLC2A3, INSIG1, and MGST1), each downregulated by about 55% in shHuR cells relative to shControl cells under non-treated conditions (Figure 1D and Table S3). One gene was significantly upregulated in shHuR relative to shControl lines, HSPA1B. For all genes SDE, there was no correlation between the fold change in expression we observed and the number of total HuR-specific PAR-CLIP conversion events [Table S5 ^17^]. Thus these changes in regulation of Nrf2-controlled transcripts are not dependent on the proclivity of the SDE transcripts to bind HuR. From this analysis, we conclude that the consensus effects, at least, are likely attributable to changes in Nrf2 regulation, as we show further below. Taken together, these data give good initial evidence that HuR is a context-dependent regulator of Nrf2 activity, modulating different subsets of Nrf2-driven genes in electrophile-stimulated vs. non-stimulated conditions.

### HuR-depleted cells manifest altered HNE-promoted antioxidant responsivity

These interesting findings, which overall are consistent with the poorly understood context dependence of Nrf2-signaling^40^, led us to examine HuR-dependent regulation of Nrf2 activity in greater detail. Specifically, we examined the extent to which cellular Nrf2-responsivity to HNE is HuR-dependent. Nrf2 activity was measured using a luciferase reporter under the transcriptional control of a Nrf2-activatable promoter, normalized to a constitutively expressed *Renilla* luciferase control (Figure 1E). Interestingly, shHuR cells under non-RES-stimulated conditions consistently featured a significant two-fold suppression of Nrf2 activity compared to shControl cells (Figure 1F), consistent with suppression of some Nrf2-driven genes revealed by our RNA-sequencing analysis. Upon ectopic expression of HuR (optimized to give only ~2.5-fold above endogenous HuR levels, Figure S4) in these shHuR cells, normal basal Nrf2 activity levels were restored (Figure 1G).

Whole-cell stimulation with HNE (25 μM; 18 h) resulted in 2-fold higher Nrf2-activity levels in HuR-depleted cells compared to shControl cells (Figure 1H). Under the same treatment conditions, ectopic expression of HuR in shHuR cells restored the extent of HNE-stimulated Nrf2-activity upregulation to that observed in shControl cells (Figure 1I). Thus, the suppression of Nrf2-activity in non-stimulated shHuR cells, as well as their greater HNE-induced Nrf2-activity upregulation observed in both our reporter assay (Figures 1F and H) and our RNA-seq data (Figures 1A and D) are HuR-specific effects.

We validated these findings in mouse embryonic endothelial cells. When these cells were depleted of HuR (Figure S5A), they also featured suppressed Nrf2 activity in the non-stimulated state (Figure S5B), and greater Nrf2 activity upregulation following HNE-stimulation [25μM; 18 h (Figure S5C)]. Consistent with our RNA-seq data, no significant difference in Nrf2 activity was observed between shHuR and shControl HEK293T cells following H_2_O_2_ stimulation (Figure 1J).

### HuR modulation of antioxidant responsivity is Nrf2-dependent

To confirm that the HuR-associated HNE-responsive effects observed above are Nrf2-dependent, we used siRNA (Figure S1B and Table S6) to simultaneously knockdown HuR and Nrf2. Multiple Nrf2 siRNAs led to ~40% suppression of Nrf2 activity on average (Figure S6A), validating the ability of our reporter assay to measure changes in Nrf2 activity. This result is also consistent with the generally narrow dynamic range of Nrf2 activity assays, as we explain further below. Knockdown of either HuR or Nrf2 in non-stimulated cells suppressed Nrf2 activity by about 50% compared to control cells; simultaneous knockdown of HuR and Nrf2 did not lead to any further suppression (Figure S6B). Upon HNE treatment, Nrf2 depletion completely arrested the ability of cells to upregulate Nrf2 activity, in the presence or absence of simultaneous HuR-knockdown (Figure S6C). Thus, the effects of HuR knockdown on Nrf2 signaling measured above are Nrf2-dependent.

The Nrf2-dependence of this HuR-regulatory event is an important finding, especially given the emerging success and promise in therapeutic targeting of Nrf2-dependent pathways^41^. However, the multifaceted regulatory modalities of Nrf2 remain poorly understood, and Nrf2-regulation at the post-transcriptional level remains an untapped arena. Thus, we chose to delve deeper—through the following series of experiments—to understand this novel post-transcriptional mechanism of Nrf2-mRNA regulation by HuR in non-stimulated cells.

### HuR and AUF1 co-regulate Nrf2 activity in non-stimulated cells

Considering that AUF1 is generally considered to be the antipode of HuR in regulating target transcripts, and that AUF1 binds to similar AU-rich elements within target transcripts as HuR, we probed the effects of both HuR and AUF1 on Nrf2 activity. We generated multiple HEK293T shAUF1 lines [each expressing a single shRNA-sequence targeting AUF1 mRNA (Table S2)] to deplete AUF1 levels. Because there are four isoforms of human AUF1 (AUF1^p37^, AUF1^p40^, AUF1^p42^, and AUF1^p45^), shRNAs targeting sequences conserved across all the isoforms were used (Table S2). Relative to a non-targeted shControl, total AUF1 was knocked down by 50–75% (Figure S1C). Because AUF1 typically destabilizes target transcripts, we expected that AUF1-knockdown would promote Nrf2 activity^20,35^. Contrary to this prediction, knockdown of AUF1 suppressed Nrf2 activity by 50-60% (Figure 2A).

**Figure 2.**
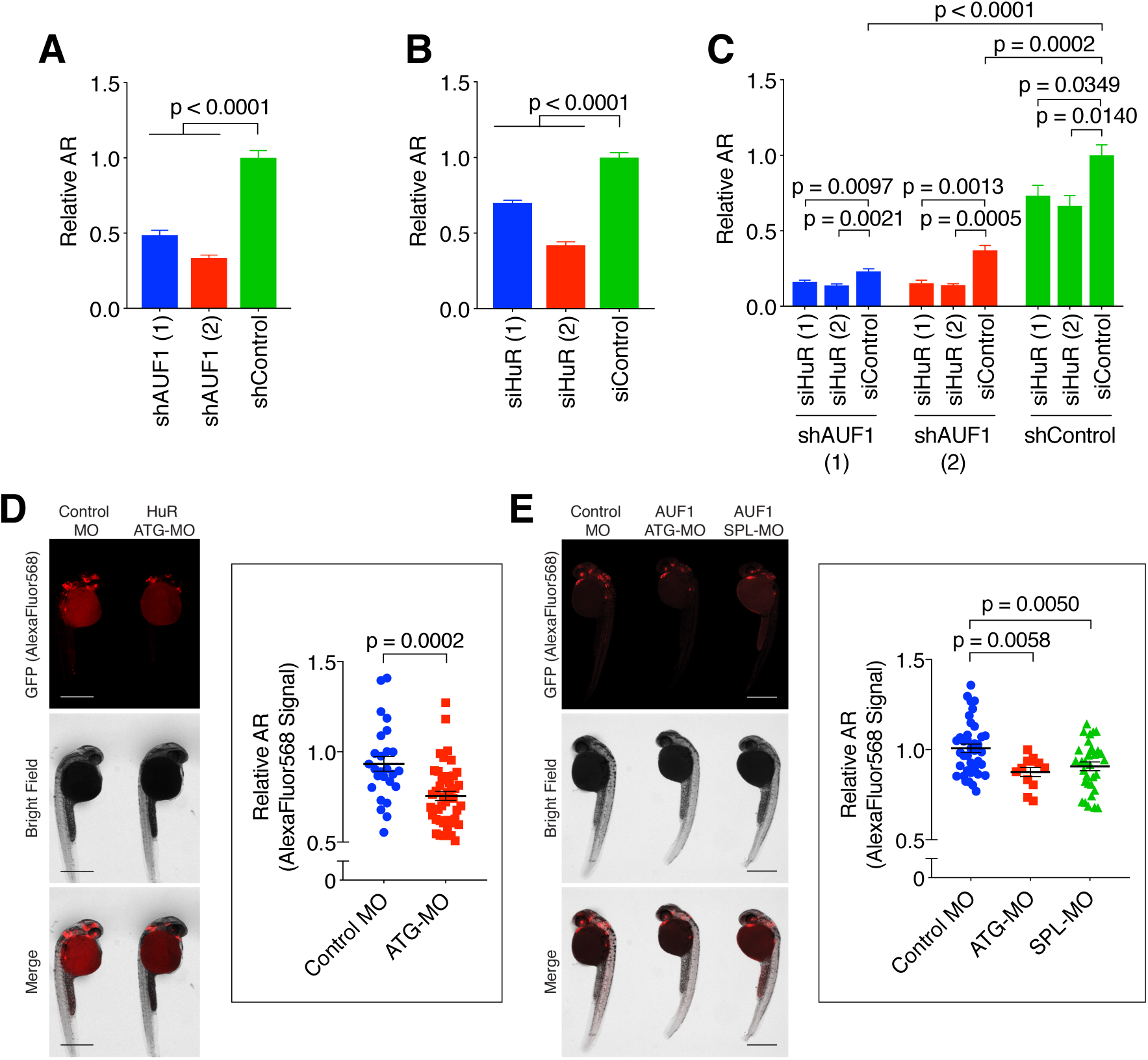
Depletion of HuR or AUF1 in cells and larval zebrafish suppresses Nrf2 activity. (**A** and **B**) Nrf2 activity in HEK293T cells depleted of HuR and AUF1, respectively [mean±SEM of n=8 (HuR) and n=12 (AUF1) independent replicates per condition]. See also Figures S1B and C. (**C**) Nrf2 activity upon simultaneous knockdown of HuR and AUF1 (mean±SEM of n=4 independent replicates). See also Figure S7A. (**D and E**) Nrf2 activity in *Tg(gstp1:GFP)* zebrafish upon knockdown of zHur and zAuf1. Larvae are stained with a red fluorescent antibody because green background fluorescence at this developmental stage prevents accurate quantitation of the GFP reporter signal. *Inset:* quantitation (mean±SEM) of mean fluorescence intensity measured using the Measure tool of ImageJ [sample sizes analyzed: E: n=26 (Control MO), 44 (ATG-MO); F: n=38 (Control MO), 12 (ATG-MO), 32 (SPL-MO)]. Each point represents a single fish. p-values were calculated with Student’s T-test. Scale bars, 500 μm. See also Figures S1D and E, and S8.

We investigated the effects of simultaneous knockdown of HuR, using the siRNAs from above, and AUF1, achieved by shRNA. Cells transfected with siHuR alone featured 40–50% suppression of Nrf2 activity (Figure 2B) relative to cells transfected with siControl. Fold suppression of Nrf2 activity upon HuR knockdown in shAUF1 (1) cells (30% on average) was not significantly different from fold suppression of Nrf2 activity upon HuR knockdown in shControl cells (Figures 2C and S7A). However, increased suppression of Nrf2 activity [relative to that observed in shAUF1 (1) and shControl cells] occurred in shAUF1 (2) (Figures 2C and S7A). These observations likely indicate that HuR- and AUF1-regulation of Nrf2 activity function through independent mechanisms. However, because Nrf2/AR has multiple upstream effectors in addition to HuR/AUF1, we further address the issue of functional (in)dependence more directly below.

### Modulation of Nrf2 activity by HuR and AUF1 is relevant at the organismal level

To extend the relevance of our findings, we turned to zebrafish (*Danio rerio*). HuR and AUF1 are both conserved in zebrafish (zebrafish gene names *elavl1a* and *hnrnpd*, respectively; we refer to these genes below as zHur and zAuf1, respectively, for clarity). The key regulators of Nrf2 pathway are also conserved in zebrafish^1^. We used antisense morpholino oligonucleotides (MOs) to deplete zHur and zAuf1 in developing embryos (Table S7) of the established Nrf2 activity-reporter strain, *Tg(gstp1:GFP)*^42^. An MO blocking translation initiation of zHur ^43^ successfully depleted zHur levels at 24 hours post-fertilization (hpf) by ~25% (Figure S1D). This is likely an underestimate of knockdown efficiency given that the commercially-available HuR antibody detects other Hu-family proteins *not* targeted by this MO. zAuf1 has only two isoforms. We designed two MOs to deplete zAuf1: one to block translation initiation (ATG-MO), and one to block a splice site (SPL-MO). Interestingly, each MO depleted only one isoform of zAuf1 (Figure S1E).

Larvae heterozygotic for the Nrf2-activity reporter were injected with MOs at the single-cell stage. 24-hours following MO-injection, we used immunofluorescent (IF) staining to assess GFP protein levels. Knockdown of either zHur or zAuf1 (by either MO for zAuf1) led to a significant 10–15% decrease in reporter protein fluorescence (Figure 2D and E). Although the magnitude of this suppression is modest, the dynamic range of Nrf2-responsivity in these fish (like many other such systems) is narrow: knockdown of *nfe2l2a* [the zebrafish Nrf2 homolog which drives AR^44^] led to only a 20%-suppression of GFP levels in these fish (Figure S8). Therefore, these data are consistent with our initial cell-based data (Figures 2A and B) and confirm that Nrf2 activity suppression promoted by depletion of HuR or AUF1 is functionally relevant in a whole vertebrate animal.

### HuR and AUF1 bind directly to Nrf2-mRNA in cells

We next confirmed that HuR/AUF1 and Nrf2-mRNA interact directly. RIP experiments were performed using HEK293T cells ectopically expressing either Flag-HuR or Flag-AUF1 isoforms. qRT-PCR analysis of eluted mRNA revealed that Nrf2 mRNA co-eluted with both HuR and all AUF1 isoforms (Figure 3A and B), confirming that both HuR and AUF1 bind directly to Nrf2-mRNA in cells.

**Figure 3.**
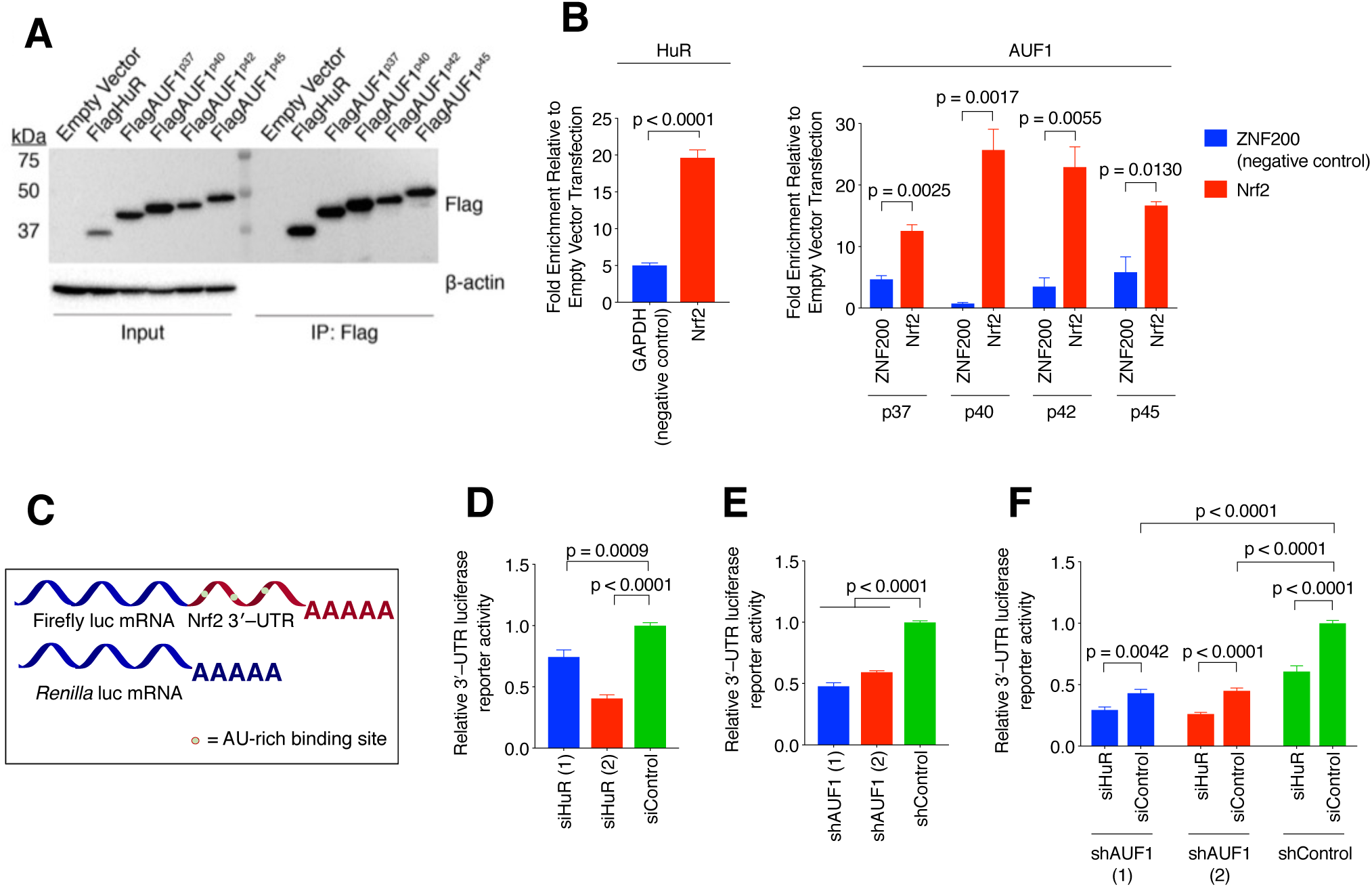
HuR and AUF1 bind directly to the 3ʹ–UTR of Nrf2-mRNA in cells. (**A**) RNA-binding protein immunoprecipitation (RIP) was carried out by expressing flag-tagged HuR or AUF1 in HEK293T cells and subjecting lysates to Flag IP. (**B**) Nrf2-mRNA co-eluting with enriched proteins was detected with qRT-PCR (mean±SEM of n=4 independent replicates). (**C**) 3ʹ–UTR reporter system consisting of Nrf2-mRNA fused to a firefly luciferase transcript and normalized to *Renilla* luciferase. (**D** and **E**) Knockdown of HuR and AUF1, respectively, reduces the 3ʹ–UTR reporter activity in HEK293T cells (mean±SEM of n=8 independent replicates for each bar). (**F**) Knockdown of both proteins leads to a further suppression of the 3ʹ–UTR reporter activity (mean±SEM of n=4 independent replicates). See also Figure S7B. All p-values were calculated with Student’s t-test.

### HuR and AUF1 bind common sites in the Nrf2 3ʹ–UTR with nanomolar affinities

The interaction of HuR and AUF1 with Nrf2-mRNA was further characterized *in vitro* by electrophoretic mobility gel shift assay (EMSA). Since HuR and AUF1 most commonly bind to sites within UTRs of target transcripts, three candidate binding sites within the 3ʹ–UTR of Nrf2-mRNA were selected (Figure S9A and Table S8) based on reported consensus binding motifs of HuR and AUF1^34,45^. Both HuR and AUF1^p37^ bound most of these sites with nanomolar affinity (Figure S9B–C; Table 1), with the exception of Nrf2 site-2, which showed negligible binding to HuR. These affinities are on par with those previously reported for HuR^46^ and AUF1^47^ to their target mRNA-binding sites, implying that the interactions with Nrf2-transcript we characterized are physiologically relevant (Table S1).

**Table 1.**
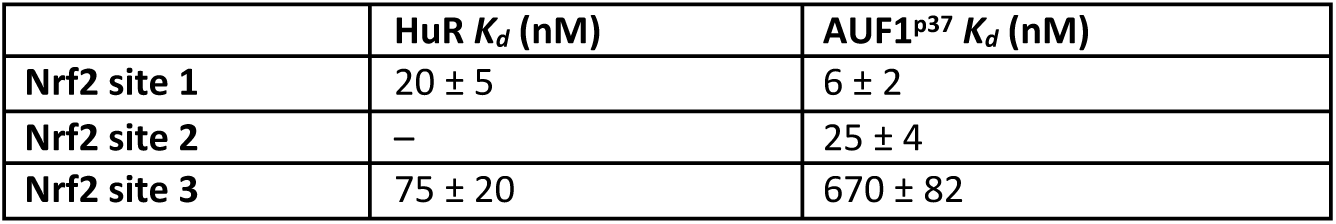
Dissociation constants measured for HuR and AUF1^p37^ binding to Nrf2 3ʹ–UTR sites.

|  | **HuR $K_d$ (nM)** | **AUF1<sup>p37</sup> $K_d$ (nM)** |
| --- | --- | --- |
| **Nrf2 site 1** | 20 ± 5 | 6 ± 2 |
| **Nrf2 site 2** | – | 25 ± 4 |
| **Nrf2 site 3** | 75 ± 20 | 670 ± 82 |

### Regulation depends on the 3ʹ–UTR of Nrf2-mRNA

At this juncture, we hypothesized that suppression of Nrf2 activity in HuR-deficient cultured cells and larval fish stems at least in part from loss of regulation at the 3ʹ–UTR of Nrf2-mRNA. To test this hypothesis, we utilized a luciferase reporter in which the 3ʹ–UTR of Nrf2 is fused to a firefly luciferase transcript^48^. A constitutively expressed *Renilla* luciferase was used as an internal normalization control (Figure 3C).

Cells deficient of HuR showed significantly reduced luciferase activity (30–60%) compared to controls (Figures 3D). We found a similar degree (40–50%) of reporter suppression in shAUF1 cells (Figure 3E). The magnitude of these effects on the 3ʹ–UTR reporter (~50%) is similar to the extent of Nrf2 activity suppression we observed in both HuR-(~50%) and AUF1-knockdown cells (~60%; Figures 2A and B). Thus, regulation of the Nrf2 3ʹ–UTR makes a significant contribution to the mechanism through which knockdown of AUF1 and HuR affects Nrf2 activity. Importantly, these findings further substantiate that the effects of HuR/AUF1-knockdown on Nrf2 activity occur specifically at the Nrf2-mRNA-level, ruling out effects of HuR/AUF1 on protein-level regulation of Nrf2.

Simultaneous knockdown of AUF1 and HuR resulted in a further decrease in luciferase activity relative to AUF1-knockdown alone (Figure 3F). Specifically, HuR knockdown suppressed 3ʹ–UTR reporter levels by ~45% regardless of the presence or absence of AUF1 (Figure S7B). Because this assay is a more direct readout of Nrf2-mRNA regulation by HuR than our Nrf2 activity reporter assay above (Figure 2C and S7A), we conclude that HuR and AUF1 act independently on Nrf2-mRNA.

### Nrf2-mRNA stability is reduced by AUF1 knockdown, but not HuR knockdown

Having established that HuR and AUF1 bind directly to Nrf2-mRNA and modulate Nrf2 activity, we sought to understand the specific mechanisms underlying this novel regulatory event. A common mode of HuR/AUF1 regulation involves stabilization or destabilization of the target upon binding^20^.

We began by considering that HuR positively regulates Nrf2 transcript stability. Levels of endogenous, mature Nrf2-mRNA were reduced in shHuR cells (20–25% relative to shControl cells Figure S10A). mRNA levels of the 3ʹ–UTR reporter transcripts were also significantly suppressed by 55% in siHuR (2)-treated cells (Figure S10B), which featured a stronger suppression in the luciferase reporter assay than siHuR (1) (Figure 3D). We expected that the half-life of Nrf2-mRNA in these cells would be similarly reduced. However, endogenous mature Nrf2 mRNA showed a non-significant (20% with 25% error) difference in half-life between shHuR and shControl cells [*t*_1/2_ = 98 (*k* = 0.007 ± 0.002 min^−1^), and 112 min h (*k* = 0.006 ± 0.0005 min^−1^), respectively (Figure S10C), which, regardless of statistical significance, is not sufficient in magnitude to explain the two-fold suppression in Nrf2 activity we consistently observed above.

Recent transcriptome-wide analysis of the stability of AUF1-targeted transcripts revealed the ability of AUF1 to positively regulate about 25% of target transcripts. Because our data showed that Nrf2 activity is suppressed in shAUF1 cells, we hypothesized that the stability of Nrf2-mRNA would be reduced in these cells. Consistent with this hypothesis, the half-life of Nrf2-mRNA was reduced by 20–60% upon knockdown of AUF1 (Figure S10D). These data also substantiate that our half-life measurements are sufficiently robust to reliably measure 2-fold changes in the stability of Nrf2-mRNA. The average suppression in half-life (50%) in shAUF1 cells is sufficient to explain the majority of the effect of AUF1-knockdown on Nrf2 activity. Thus, AUF1 *positively* regulates Nrf2 activity by stabilizing Nrf2-mRNA. Collectively, the mechanistic differences between the effect of AUF1 and HuR (which we elaborate on below) further point to independent effects of HuR and AUF1 on Nrf2 activity.

### HuR enhances Nrf2-transcript splicing

Having ruled out significant effects on Nrf2-mRNA stability upon HuR knockdown, we investigated other potential means by which HuR regulates Nrf2 activity. In addition to its classical role in stabilizing target mRNAs, HuR also regulates maturation of some of its targets^23^. We reasoned that modulations in Nrf2-mRNA processing (e.g. splicing, translocation) could lead to decreased levels of mature Nrf2-mRNA and contribute to the suppression in Nrf2 activity we observed. Importantly, premature Nrf2-mRNA does not contribute to the pool of functional Nrf2-mRNA (competent for translation into Nrf2-protein), because it is not exported from the nucleus and cannot be translated to give a functional protein (Figure S11A).

We first turned to our RNA-seq data for evidence of misregulation of maturation. Because we prepared our RNA-sequencing library using rRNA depletion and deep-sequenced the resulting library, we were able to examine the levels of intronic RNA present in the Nrf2 transcript. Nrf2-mRNA is composed of 5 exons with 4 intervening introns. The first intron of Nrf2 is extremely long at ~30,000 bp (Figure S11B), putting it in the longest 10% of human introns^49^. When we examined the levels of intronic RNA in the Nrf2-transcript in shHuR and shControl cells, we found that Nrf2 transcripts in shHuR cells contained higher levels of intronic content (Figure 4A). Thus, knockdown of HuR *suppresses* the maturation of Nrf2-mRNA, leading to an accumulation of unspliced, premature Nrf2-mRNA.

**Figure 4.**
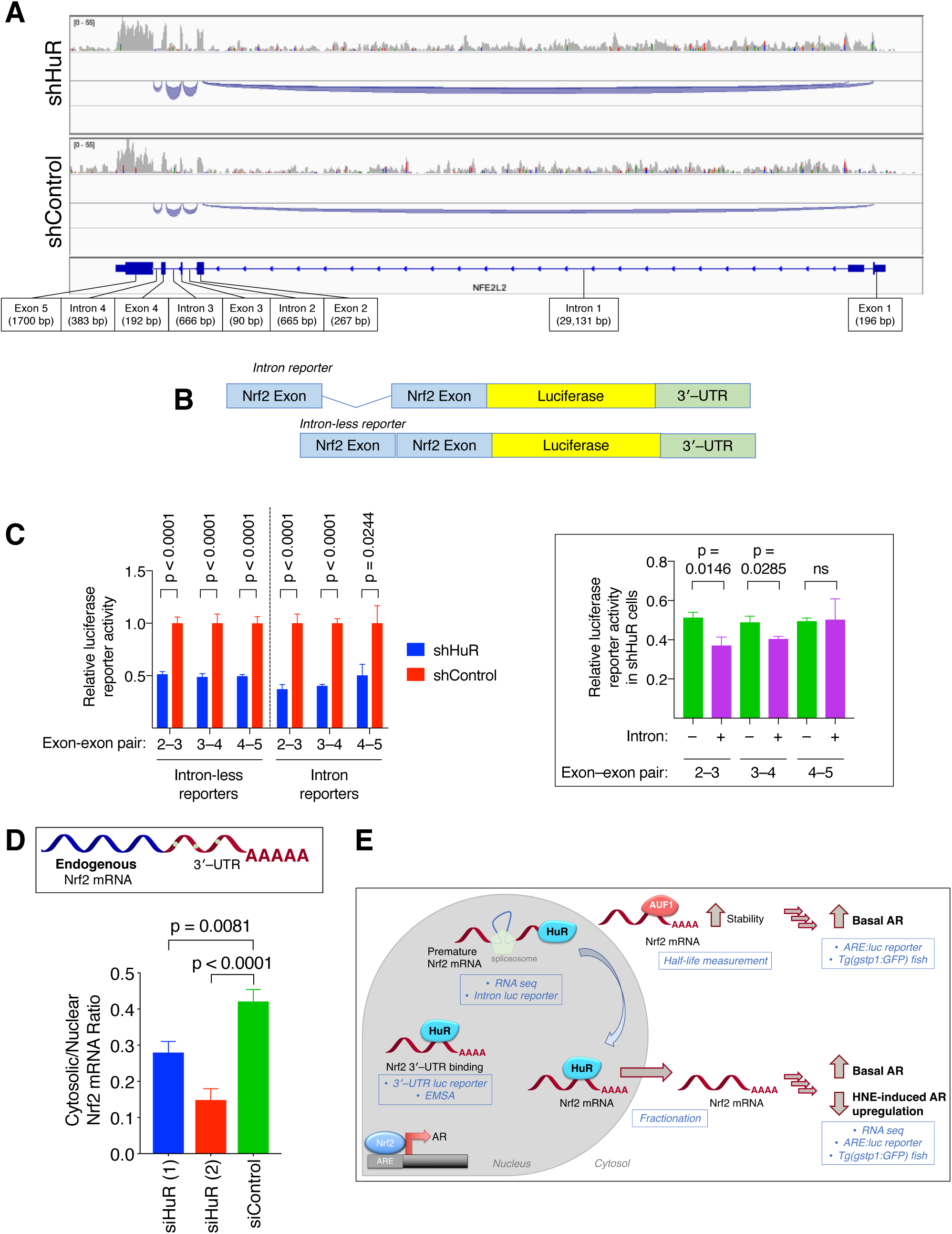
HuR-knockdown suppresses Nrf2-mRNA splicing and nuclear export of Nrf2-mRNA. (**A**) Integrative Genomics Viewer ^70^ view of splicing tracks for Nrf2 (NFE2L2) from RNA-sequencing analysis of shHuR and shControl HEK293T cells. The area of the blue tracks corresponds to the levels of intronic RNA detected. Shown is one representative replicate per cell line. See also Figure S11. (**B**) Reporters to readout the effect of introns were constructed by fusing a portion of Nrf2-mRNA (with or without the intron) upstream of a firefly luciferase reporter. As with the 3ʹ–UTR reporter (Figure 3C), effects on this construct can be assayed by measuring luciferase activity in the lysates of cells expressing these reporters. (**C**) Reporter levels upon HuR knockdown in non-stimulated HEK293T cells for both the intron-less reporter (left) and the intron reporter (right) (mean±SEM of n≥7 per set). *Inset at right*: Comparison of reporter activity in shHuR cells (i.e., blue bars from the main plot in **C** upon introduction of introns. (**D**) qRT-PCR was used to measure the ratio of endogenous Nrf2-mRNA in nuclear and cytosolic extracts of HEK293T cells depleted of HuR (mean±SEM of n=8 for siHuR (1 and 2) and n=7 for siControl). (**E**) Model of posttranscriptional regulation of Nrf2-mRNA by HuR and AUF1. HuR regulates Nrf2-mRNA maturation and nuclear export, and AUF1 stabilizes Nrf2-mRNA. Shown in blue text/boxes are the experimental evidence supporting each facet of this regulatory program. All p-values were calculated with Student’s t-test.

To probe this effect further, we utilized a firefly luciferase reporter with either an exon–intron–exon (“intron reporter”) sequence or an exon–exon (“intron-less reporter”) sequence from Nrf2-mRNA fused upstream of the luciferase reporter. The Nrf2 3ʹ–UTR was also retained downstream of the luciferase reporter. Reporters were constructed containing 3 different Nrf2 exon pairs, with or without the intervening introns (Figure 4B). Regulation of these mRNA reporters can be assessed by measuring firefly luciferase enzyme activity in lysates of cells expressing these reporters (normalized to a constitutively expressed *Renilla* luciferase control). Because of the length of the 1^st^ intron of Nrf2 (Figure S11B), this intron was not investigated in this assay.

Expression of the intron-less reporters in shHuR cells led to a 50% decrease in reporter activity relative to shControl cells (Figure 4C), consistent with our 3ʹ–UTR luciferase reporter data above (Figure 3D). Because this effect (~50% reduction in shHuR cells) was similar across all three reporters containing different exons, it is likely attributable to the 3ʹ–UTR (the common factor between the three reporters). However, two of the intron-containing reporters showed a further suppression upon HuR knockdown relative to their intron-less counterparts (Figure 4C, inset at right), consistent with our RNA seq data (Figure 4A). The greatest suppression effects were observed for introns 2 and 3, whereas intron 4 was not strongly suppressed in shHuR cells in either the RNA-seq or the reporter data.

Overall, these data agree with PAR-CLIP results indicating nucleotide conversions (i.e., HuR binding) within Nrf2 introns, as well as the presence of multiple putative binding sites in the introns based on consensus HuR-binding sequences^17^. These data collectively indicate that HuR enhances the splicing and maturation of Nrf2-mRNA. The magnitude of this effect is on par to the suppression of endogenous Nrf2-mRNA levels we observed above.

### HuR regulates nuclear export of Nrf2-mRNA

Nuclear export of mature mRNA is intrinsically linked to the splicing/maturation process because maturation is a prerequisite for export. Coupling of these processes is an appreciated mechanism to enhance gene expression^50^. mRNA that is not exported due to improper or incomplete splicing is rapidly degraded in the nucleus^51^. Given the ability of HuR to shuttle between the nucleus and cytosol^21^, we hypothesized that HuR governs Nrf2-mRNA export from the nucleus. Indeed, for some mRNA targets such as COX-2, HuR plays a key role in regulating export from the nucleus^22^. To test whether a similar regulatory program is at play for Nrf2-mRNA, we employed nuclear/cytosolic fractionation in shHuR/shControl cells coupled with qPCR detection of endogenous, mature Nrf2-mRNA. These experiments revealed that the cytosolic/nuclear ratio of endogenous Nrf2-mRNA decreased by 35% on average in HuR-deficient cells (Figure 4D), indicating a *decrease in Nrf2-mRNA nuclear export* upon depletion of HuR. Faster turnover of nuclear-mRNA may contribute to the reduction in overall levels of Nrf2-mRNA upon HuR-knockdown (Figure S10A) and to the slight reduction in Nrf2-mRNA half-life we observed (Figure S10C). In sum, we conclude that the principal mechanisms by which HuR regulates Nrf2-mRNA are control of Nrf2-mRNA splicing/maturation and nuclear export of the mature transcript. Importantly, these results collectively point to a novel regulatory program that functions specifically at the Nrf2-mRNA level to modulate Nrf2 activity (Figure 4E).

## DISCUSSION

We began by exploring HuR-dependent global transcriptomic changes in cells stimulated with RES/ROS, using H_2_O_2_ and HNE as representative native ROS/RES. Contrary to previous reports of the sensitivity of HuR to oxidative stress^52^, we found no transcriptional responses specifically attributable to HuR following H_2_O_2_ stimulation (Figure 1B). Differences in cell type may explain this observed discrepancy, as we observed a muted transcriptional response to H_2_O_2_ stimulation overall in our HEK293T cell lines. Conversely, HNE-stimulation of HuR-knockdown cells elicited a general downregulation of global transcriptional activity compared to control cells (Figure 1A and S3). Given the important roles that pleiotropic covalent drugs and native RES play on Nrf2 signaling^2,3^, we homed in on differentially-expressed Nrf2-driven genes. This analysis revealed an interesting bifurcation between HuR-dependent effects on Nrf2-driven genes in non-stimulated shHuR cells versus HNE-stimulated cells. Nrf2-driven genes that were more strongly upregulated in HNE-stimulated shHuR cells (ME-1, TXNRD1, and FTL) were not significantly differentially expressed in non-stimulated shHuR cells relative to shControl cells (Figures 1A and C). However, a separate subset of Nrf2-driven genes (SLC2A3, INSIG1, and MGST1) was suppressed in non-stimulated shHuR cells (Figure 1D). Together with our Nrf2 activity reporter data (Figures 1F and H) which was consistent with the majority of our RNA seq analysis, these findings highlight context-specific complexities inherent in the regulation of Nrf2 pathway by HuR that echo recent reports in the field^40^. Because of these intricacies, we chose to focus our further investigations on non-stimulated cells.

For both HuR and AUF1, which we also identified as a novel regulator of Nrf2 activity, we found 2–3-fold suppression in Nrf2 activity in both HuR- and AUF1-knockdown cells relative to control cells (Figures 1F, 2A, 2B, and S5). These changes are indicative of significant regulatory events because we and others have previously documented the narrow dynamic range (2–4-fold) of Nrf2 activity modulation in various readouts, including luciferase reporter assays, flow cytometry-based analyses, as well as qRT-PCR and western blot analyses which assay endogenous Nrf2-driven genes^53–56^. Consistent with this logic, knockdown of Nrf2 gave a similar fold change in Nrf2 activity to HuR knockdown. There was no further decrease in Nrf2 activity when HuR and Nrf2 were simultaneously knocked down, clearly showing that the fold change observed in Nrf2 activity is the maximum possible in the system, and that HuR functions through modulating Nrf2 activity. Two-to three-fold changes are also typical magnitudes of response in other pathways regulated by electrophiles. For instance, knockdown of the electrophile sensor Pin1 elicits only about 30% decrease in viability upon treatment with HNE relative to knockdown control^57^. Our findings that HuR knockdown-induced Nrf2 activity suppression extends to whole zebrafish depleted of HuR or AUF1 (Figures 2D and E), as well as to mouse endothelial cells MEECs (Figure S5B) demonstrate a broader generality that is consistent with the fact that HuR and AUF1 are ubiquitously expressed. These novel general regulatory roles of HuR/AUF1 on Nrf2 mRNA stand in contrast to previously reported Nrf2-mRNA regulatory events by miRNAs that are highly cell/context specific^8^. Thus, in terms of cell type generality and magnitude, our study goes a long way in demonstrating that Nrf2-mRNA regulation is an important modulator of Nrf2 activity.

HuR and AUF1 can function in concert to destabilize mRNAs such as p16^INK4 58^, and enhance the translation of mRNAs such as TOP2A^34^, among other examples. However, we found that HuR and AUF1 act independently on Nrf2-mRNA, despite similar binding affinities to common sites we identified within the 3ʹ–UTR (Table 1). The distinct, orthogonal mechanisms by which HuR and AUF1 regulate Nrf2-mRNA— control of Nrf2-mRNA splicing and export; and stabilization of Nrf2-mRNA, respectively—agree with our Nrf2 activity reporter data indicating independent effects of HuR and AUF1 on Nrf2/AR-axis. These effects could not have been predicted based on the previously-reported PAR-CLIP binding data alone. The complexities we uncovered in this system collectively speak to the need for careful mechanistic evaluation of “on-target” effects of HuR/AUF1-knockdown, particularly when common binding sites are involved.

As is appreciated in the mRBP field^17,35^, both HuR and AUF1 regulate their targets through multiple mechanisms. Although the magnitude of the UTR-reporter suppression (Figure 3D) and nuclear-export suppression (Figure 4E) we observed in HuR-knockdown cells at first glance seem sufficient to explain the magnitude of Nrf2 activity suppression, our data testify that additional regulatory modes beyond UTR-regulation are at play. For instance, using multiple approaches, we identified Nrf2-mRNA splicing as a subtle but significant component of HuR regulation of Nrf2-mRNA (Figure 4A–C). These alternate/complementary mechanisms may explain why we unexpectedly observed a reduction in the global pool of Nrf2-mRNA without a similar fold-change in the half-life of Nrf2-mRNA upon HuR-knockdown, for instance. Our mechanistic interrogations ultimately established that reduced splicing and reduced nuclear export of Nrf2-mRNA are the principal mechanisms by which Nrf2 activity is suppressed upon HuR-knockdown. These mechanisms are likely linked as splicing is generally a prerequisite for nuclear export^50^, although the details of this mechanistic coupling for Nrf2-mRNA remain to be explored. Nevertheless, downregulation of both splicing/maturation and nuclear export of Nrf2-mRNA upon HuR-knockdown likely explains why we observed a suppression in the pool of mature Nrf2-mRNA (Figure S10A). Overall, the mechanisms we identified strongly and clearly point to on-target, Nrf2-mRNA-specific events that support HuR/AUF1 regulation of Nrf2 signaling.

In sum, these data highlight an important intersection between proven disease-relevant players: HuR, AUF1, and Nrf2 are all upregulated in cancers^59,60^; therapeutic targeting of Nrf2/AR axis through exploitation of protein regulators such as Keap1 continues to be a promising small-molecule intervention^7,61^. The ubiquitous expression of HuR and AUF1, along with the conservation of their regulation of Nrf2 activity across multiple cell types and whole organisms underscore the importance of this newly identified regulatory program. Because this regulation of Nrf2/AR occurs specifically at the mRNA-level, it potentially offers an orthogonal therapeutic strategy to modulate this conserved pathway of validated pharmacological significance.

## Materials and Methods

### Statistics and data presentation

For experiments involving cultured cells, samples generated from individual wells or plates were considered biological replicates. For zebrafish experiments, each larval fish was considered a biological replicate. In the figure legend for each experiment, the number of independent biological replicates and how the data are presented in the figure (typically mean ± SEM) are clearly indicated. P-values calculated with Student’s unpaired t-test are clearly indicated within data figures. Data were plotted/fit and statistics generated using GraphPad Prism 7 or 8.

### Reagents

All HNE used in this study was HNE(alkyne) (referred to as HNE in the manuscript/figures for clarity), and was synthesized as previously reported ^54^. Unless otherwise indicated, all other chemical reagents were bought from Sigma at the highest availability purity. TCEP was from ChemImpex. Puromycin was from Santa Cruz. Actinomycin D was from Sigma. AlamarBlue was from Invitrogen, and was used according to the manufacturer’s instructions. Minimal Essential Media, RPMI, Opti-MEM, Dulbecco’s PBS, 100X pyruvate (100 mM), 100X nonessential amino acids (11140-050) and 100X penicillin streptomycin (15140-122) were from Gibco. Protease inhibitor cocktail complete EDTA-free was from Roche. 3XFlag peptide was from APExBIO. Anti-Flag(M2) resin (A2220) was from Sigma-Aldrich. TALON (635503) resin was from Clontech. Ni-NTA agarose (30210) was from QIAGEN. 2020 and LT1 transfection reagents were from Mirus. DharmaFECT I and Duo were from Dharmacon. PEI was from Polysciences. Venor GeM PCR-based mycoplasma detection kit was from Sigma. ECL substrate and ECL-Plus substrate were from Pierce and were used as directed. Acrylamide, ammonium persulfate, TMEDA, Precision Plus protein standard were from Bio-Rad. All lysates were quantified using the Bio-Rad Protein Assay (Bio-Rad) relative to BSA as a standard (Bio-Rad). PCR was carried out using Phusion Hot start II (Thermo Scientific) as per the manufacturer’s protocol. All plasmid inserts were validated by sequencing at Cornell Biotechnology sequencing core facility. All sterile cell culture plasticware was from CellTreat.

### Generation of plasmids

Sequences of all primers used for cloning and site-directed mutagenesis are listed in Table S9. All plasmids generated were fully validated by Sanger sequencing at the Cornell Genomics Core Facility.

pCS2+8 Flag_3_HuR was generated by PCR-amplification of human HuR (plasmid provided from the Hla lab), extension of the resulting product, and cloning into linearized pCS2+8 vector (Addgene plasmid #34931). For recombinant expression, the same procedure was used to clone HuR into a pET28a vector.

pLJM60 AUF1^p42^ was obtained from Addgene (plasmid #38242) and cloned with a Flag_2_ tag into pCS2+8 as described above. Using the resulting plasmid, AUF1^p37^ and AUF1^p45^ were generated by deletion of AUF1-exon 7 and addition of AUF1-exon 2, respectively. AUF1^p40^ was generated by addition of AUF1-exon 7 to AUF1^p37^. For recombinant expression, these constructs were cloned into a pET28a vector.

The pSGG luciferase reporter plasmid ^48^ containing the Nrf2 3ʹ–UTR was obtained from Prof. Qun Zhou (University of Maryland School of Medicine). This plasmid was modified for intron-reporter assays by cloning fragments of the Nrf2 transcript with or without introns (amplified from genomic DNA or cDNA prepared from HEK293T cells, respectively) upstream of the luciferase coding sequence.

shRNA-resistant expression plasmids (for rescue experiments) were produced by PCR amplification of the starting plasmid with forward and reverse mutagenesis primers containing the desired mutations (Table S9) followed by DpnI (NEB) treatment. These plasmids code for the same protein but contain mismatches (highlighted in red in the primer sequences; Table S9) in the shRNA-targeting sequence allowing expression of the ectopic protein in knockdown cells.

### Cell culture

HEK293T (obtained from ATCC) and MEECs (generated in the Hla lab) were cultured in MEM (Gibco 51090036) supplemented with 10% v/v fetal bovine serum (Sigma), penicillin/streptomycin (Gibco), sodium pyruvate (Gibco), and non-essential amino acids (Gibco) at 37°C in a humidified atmosphere of 5% CO_2_. Media was changed every 2-3 days.

### Generation of shRNA-based knockdown cell lines

HEK293T packaging cells (5.5 × 10^5^ cells) were seeded in 6-well plates in antibiotic-free media and incubated for 24 h. The cells were transfected with a mixture of 500 ng packaging plasmid (pCMV-R8.74psPAX2), 50 ng of envelope plasmid (pCMV-VSV-G) and 500 ng of pLKO vector containing the hairpin sequence (Sigma; Table S2) using TransIT-LT1 (Mirus Bio) following the manufacturer’s protocol. shControl plasmid was obtained from Prof. Andrew Grimson (Cornell University). 18 h-post transfection, the media was replaced with media containing 20% FBS and incubated for a further 24 h. Media containing virus particles was harvested, centrifuged (800×g, 10 min), filtered through 0.45 μm filter, and used for infection or frozen at –80°C for later use.

HEK293T or MEEC cells (5.5 × 10^6^ cells) in 6-well plates were treated with 1 mL of virus-containing media in a total volume of 6 mL of media containing 8 μg/mL polybrene. After 24 h, media was changed and the cells were incubated for 24 h. Following this period, media was changed to media containing 2 μg/mL puromycin (Santa Cruz). Following selection, cells were assayed by western blotting.

### RNA sequencing

Following treatment of shHuR/shControl cells with HNE (50 μM) or H_2_O_2_ (225 μM) for 18 h, cells were lysed by the addition of Trizol (Invitrogen) and RNA was isolated following the manufacturer’s protocol. The quality of the RNA was assessed by Nanodrop spectrophotometry (A260/A280 ratio ~2) and fragment analysis using a Bioanalyzer (Agilent). RNA quality numbers (RQNs) for all samples were 10.0. The rest of the procedure including data analysis was performed by the RNA sequencing core (RSC) service at Cornell University. Briefly, ribosomal (r)RNA was subtracted by hybridization from total RNA samples (100 ng total RNA input) using the RiboZero Magnetic Gold H/M/R Kit (Illumina). Following cleanup by precipitation, rRNA-subtracted samples was quantified with a Qubit 2.0 (RNA HS kit; Thermo Fisher). TruSeq-barcoded RNAseq libraries were generated from half of the rRNA-subtracted samples with the NEBNext Ultra II Directional RNA Library Prep Kit (New England Biolabs). Each library was quantified with a Qubit 2.0 (dsDNA HS kit; Thermo Fisher) and the size distribution was determined with a Fragment Analyzer (Advanced Analytical) prior to pooling. Libraries were sequenced on a NextSeq500 instrument (Illumina). At least 60M single-end 75bp reads were generated per library. Reads were trimmed for low quality and adaptor sequences with cutadapt v1.8 [parameters: –m 50 –q 20 –a AGATCGGAAGAGCACACGTCTGAACTCCAGTC –match-read-wildcards ^62^]. Trimmed reads were mapped to rRNA sequences with bowtie2 v2.2 to remove them [Parameters: default mapping options, –al and – un to split matching and non-matching reads ^63^]. Reads were mapped to the reference genome/transcriptome (GRCh37/hg19]) using tophat v2.1 [parameters: –library-type=fr-firststrand –no-novel-juncs –G <ref_genes.gtf> ^64^]. Cufflinks v2.2 (cuffnorm/cuffdiff) was used to generate FPKM values and statistical analysis of differential gene expression ^39^.

### Growth inhibition assays

HEK293T cells (4000 cells per well) were seeded in 96-well plates. After 24 h, cells were treated with the indicated molecules at the indicated concentrations and incubated for 48 h. AlamarBlue (Invitrogen) was added to each well and the cells were incubated for a further 3 h, after which fluorescence (excitation 560 nm; emission 590 nm) was measured using a BioTek Cytation 3 microplate reader. To test whether reduced alamarBlue is oxidized by H_2_O_2_ over the timescales used, HEK293T cells in a 96-well plate were incubated with the Manufacturer’s recommended concentration of alamarBlue for 3 h to reduce the dye. The growth medium containing reduced alamarBlue was collected, pooled, centrifuged to remove cells, and then incubated with the concentrations of H_2_O_2_ used in the viability experiment (Figure S2B) and fluorescence was measured over time as described above. We observed ~3% loss of fluorescence after 4 h in reduced alamarBlue treated with 1250 μM H_2_O_2_; the fluorescence of samples at all other concentrations was not significantly different from equivalent media containing reduced alamarBlue but not treated with H_2_O_2_. Thus, this control experiment confirmed that our experimental conditions involving oxidants are compatible with redox-sensitive AlamarBlue-reagents employed in our growth inhibition assays.

### Knockdown of HuR and Nrf2 with siRNA

SiRNAs targeting the ORF of human HuR or Nrf2 were obtained from Dharmacon (Table S6) and non-targeting control siRNAs (Control siRNA-A or E) were obtained from Santa Cruz Biotechnology. HEK293T (3.6 × 10^5^ cells) in 6-well plates were transfected with siRNA using Dharmafect I (Dharmacon) for 48 h following the manufacturer’s protocol, then assayed. For co-transfection of siRNA and plasmid(s) for reporter assays, cells were transfected with Dharmafect Duo (Dharmacon) for 48 h following the manufacturer’s protocol, then assayed.

### Western blotting

Cells were resuspended in RIPA buffer (Santa Cruz Biotechnology) supplemented with protease and phosphatase inhibitors and lysed by three cycles of rapid freeze-thaw. Lysates were cleared by centrifugation (20000 × g, 10 min, 4°C) and total protein concentration was determined by the Bradford assay. Typically, 20–40 μg of total protein was loaded per lane, separated by SDS-PAGE, transferred to PVDF, blocked, and incubated with the appropriate antibodies (Table S10). Detection was carried out on a ChemiDoc-MP imaging system (BioRad) using ECL Western Blotting Substrate (Pierce) or SuperSignal West Femto Maximum Sensitivity Substrate (Thermo Scientific). Western blot data were quantitated using the Gel Analysis tool in ImageJ (NIH). Bands of interest were integrated and normalized to the loading control.

### Luciferase reporter assays

Luciferase reporter assays were carried out as previously described ^65^. Briefly, cells were co-transfected with firefly luciferase plasmid [pGL4.37 E364A for ARE(antioxidant response element):luciferase assays (Promega); pSGG containing the human Nrf2 3ʹ–UTR sequence (from Prof. Qun Zhou, University of Maryland School of Medicine)] and *Renilla* luciferase plasmid (pGL4.75, E693A (Promega)) in a 40:1 ratio for 48 h using TransIT 2020 (Mirus Bio) or Dharmafect Duo (Dharmacon). For experiments involving ectopic protein expression, the 40:1 reporter plasmid mixture was co-transfected with ectopic protein expression plasmid in a 1:1 ratio. Cells were lysed for 15 min in passive lysis buffer (Promega) with gentle shaking and homogenized lysate was transferred to the wells of an opaque white 96-well plate (Corning). Firefly and *Renilla* luciferase activity was measured on a BioTek Cytation3 microplate reader. For assays involving HNE treatment, media were replaced with media containing the indicated concentration of HNE at 30 h post-transfection, and cells were incubated for a further 18 h.

### Quantitative real time PCR (qRT-PCR)

qRT-PCR was carried out as previously described ^53^. Total RNA was isolated from cells using Trizol reagent (Ambion) following the manufacturer’s protocol. 1 μg of total RNA (purity/integrity assessed by agarose gel electrophoresis and concentration determined by A_260nm_ using a BioTek Cytation3 microplate reader with a Take3 accessory) was treated with amplification-grade DNase I (Invitrogen) and reverse transcribed using Oligo(dT)_20_ as a primer and Superscript III Reverse Transcriptase (Life Technologies) following the manufacturer’s protocol. PCR was performed for two technical replicates per sample using iQ SYBR Green Supermix (BioRad) and primers specific to the gene of interest (Table S11) following the manufacturer’s protocol. Amplicons were chosen that were 150–200 bp in length and had no predicted off-target binding predicted by NCBI Primer BLAST. For genes with multiple splice variants, primers were chosen that amplified conserved sequences across all splice variants. Primers were validated using standard curves generated by amplification of serially-diluted cDNA; primers with a standard curve slope between –0.8 and 1.2 and R^2^≥0.97 were considered efficient. Single PCR products were confirmed by melting analysis following the PCR protocol. Data were collected using a LightCycler 480 (Roche). Threshold cycles were determined using the LightCycler 480 software. Samples with a threshold cycle >35 or without a single, correct melting point were not included in data analysis. Normalization was carried out using a single housekeeping gene as indicated in each dataset and the ΔΔC_t_ method.

### Analysis of reporter mRNA levels

HEK283T cells (1.6 × 10^5^) in 12-well plates were transfected with luciferase reporters as described above for 48 h. Cells were then lysed in 250 μl of passive lysis buffer for 15 min with shaking and 50 μl of lysate was taken to measure luciferase activity as described above. Trizol LS was added to the remaining lysate and RNA was isolated as described above. Prior to reverse transcription, total RNA (1μg) was treated with amplification-grade DNase I (Invitrogen) to remove any residual plasmid. Reverse transcription and qRT-PCR was then carried out as above.

### RIP-PCR

RIP-PCR was carried out following previously described methods ^66^. HEK293T cells (4.5 × 10^6^) in 10 cm diameter plates were transfected with the indicated constructs (mixed 1:3 with empty plasmid) or empty plasmid alone (8 μg total DNA per plate) with PEI (21 g per plate). Media was changed 24 h-post transfection and the cells were incubated 48 h total. Cells were washed once with PBS (Invitrogen) and harvested by trypsinization, and washed thoroughly with PBS.

Cell pellets were resuspended in polysome lysis buffer [10 mM HEPES (Chem-Impex) pH 7.0, 100 mM KCl (Fisher), 5 mM MgCl_2_ (Fisher), 0.5% Nonidet-P40, cOmplete EDTA-free Protease Inhibitor Cocktail (Roche), 0.2% vanadyl ribonuceloside complex (NEB)] and frozen at –80°C for at least 30 min. The lysate was thawed, centrifuged twice (20000 × g, 10 min each), and the protein concentration was determined using Bradford Assay. 3 mg of total protein was diluted to 0.5 mg/mL in NT2 buffer [50 mM Tris pH 7.4, 150 mM NaCl (Fisher), 1 mM MgCl_2_ (Fisher), 0.05% Nonidet-P40, 0.2% vanadyl ribonuceloside complex] and incubated with 50 μL of Flag M2 agarose (Sigma) for 3 h at 4°C with end-over-end rotation. The resin was washed at least 4 times (5 min per wash) with NT2 buffer containing 300 mM NaCl and a portion of the resin was retained for western blot analysis. The resin was resuspended in 100 μL of NT2, supplemented with 0.1% SDS and 3 mg/mL proteinase K (Santa Cruz Biotechnology), and incubated at 55°C for 30 min. RNA was isolated using Trizol reagent following the manufacturer’s protocol and analyzed by qRT-PCR as described above.

### Analysis of mRNA stability

HEK293T shHuR/shAUF1 and shControl cells (9.6 × 10^5^ cells) in 6-well plates were treated with 5 μg/mL actinomycin D (Sigma) and harvested in Trizol as described above at the indicated time points. The level of Nrf2-mRNA remaining was assessed by qRT-PCR as described above.

### Nuclear-cytosolic fractionation for RNA isolation

This procedure was adapted from a reported protocol ^67^. HEK293T cells (9.6 × 10^5^ cells) in 6-well plates were harvested by tryspinization and washed twice with cold PBS. Cell pellets were then resuspended in RSB buffer [10 mM Tris (ChemImpex) pH 7.4, 10 mM NaCl (Fisher), 3 mM MgCl_2_ (Fisher)], incubated on ice for 3 min, and centrifuged (1500 × g, 3 min, 4°C). Cells were resuspended in RSBG40 buffer [10 mM Tris pH 7.4, 10 mM NaCl, 3 mM MgCl_2_, 10% glycerol (Millipore), 0.5% Nonidet P-40, 0.5 mM DTT (VWR), and 100U/ml RNaseOUT (Invitrogen)] and lysed with gentle pipetting up and down. The suspension was centrifuged (4500 × g, 3 min, 4°C) and the supernatant was saved as the cytosolic fraction. The nuclear fraction (pellet) was again resuspended in RSBG40 and 10X detergent solution [final concentrations: 3.3% sodium deoxycholate (ChemImpex), 6.6% Tween 20 (Fisher)] was added. The nuclei were pipetted up and down several times and the suspension was incubated on ice for 5 min and then centrifuged (4500 × g, 3 min, 4°C). The supernatant was pooled with the previous supernatant and the combined sample was used as the cytosolic fraction. The nuclei were washed once more with RSBG40, centrifuged (9500 × g, 3 min, 4°C), and resuspended thoroughly in RSBG40. Trizol LS was then added to each sample and RNA was isolated, treated with amplification-grade DNase I, and subjected to qPCR as described above.

### Zebrafish husbandry, microinjection, and imaging

All zebrafish procedures were performed in accordance with the guidelines of NIH and were approved by Cornell University’s Institutional Animal Care and Use Committee (IACUC). *Tg(−3.4gstp1:GFP)it416b* fish were obtained from RIKEN Brain Science Institute, Japan (National BioResource Project, Zebrafish). Transgenic fish were crossed with wild-type (WT) fish to generate a mixture of heterozygous and WT progeny for experiments. Fertilized eggs at the single-cell stage were injected with 8 ng of morpholino oligonucleotides (MOs) targeting zHuR (*elavl1a*) or zAUF1 (*hnrnpd*) or a random control MO (Gene Tools, LLC.; Table S7). Fish were screened for GFP expression by imaging with a Leica M205-FA fluorescence stereoscope approximately 24 h post-fertilization (hpf). Following imaging, fish were euthanized, washed twice in PBS with 0.1% Tween-20, and transferred to 4% formaldehyde for further IF analysis (see below). GFP expression was detected using red fluorescent antibody staining because of high background signal in the GFP (ex: 488nm; em: 520–550 nm) channel at this zebrafish developmental stage which prevents accurate quantitation.

### Whole-mount immunofluorescence

Whole-mount immunostaining of zebrafish larvae was carried out as previously described ^68^. All incubation steps were performed with gentle rocking. Euthanized fish were washed twice with PBST (PBS + 0.1% Tween-20) and fixed in 4% formaldehyde in PBS at 4°C overnight or up to 7 days. Fish were then washed twice for 30 min with PDT (PBST, 0.3% Triton-X100, 1% DMSO), blocked for 1 h at room temperature in blocking buffer (PBST, 10% FBS, 2% BSA), and incubated with primary antibody (Table S10) in blocking buffer for 2 h at room temperature or overnight at 4°C. Fish were washed twice with PDT, re-blocked for 1 h at room temperature, and incubated with secondary antibody (Table S10) in blocking buffer for 1.5 h at room temperature. Following secondary antibody incubation, fish were washed twice with PDT and then imaged. Fluorescence was quantified using the “Measure” tool of ImageJ (NIH).

### Expression and purification of His_6_-HuR

BL21-CodonPlus cells (Agilent) were transformed with pET28a plasmid encoding His_6_-TEV-HuR. A single colony was divided into several starter cultures (5 ml) and grown in LB media containing chloramphenicol and kanamycin overnight at 37°C with shaking. Cultures were then each diluted into 1 L of LB containing kanamycin and grown at 37°C with shaking to OD_600 nm_ =0.5, at which point IPTG (Gold Biotechnology; 1 mM final concentration) was added to induce expression. The temperature was then reduced to 18°C and cultures were grown overnight with shaking. Further steps were carried out at 4°C. Cells were harvested by centrifugation (4000 × g, 20 min, 4°C), resuspended in lysis buffer [20 mM Tris pH 7.5, 500 mM KCl, 2 mM MgCl_2_, 10 mM Imidazole (Fisher), 5% glycerol, 5 mM βME, and 0.5 mM PMSF (Alexis Biochemicals)], and lysed by two passages through an Emlusiflex cell disruptor (Avestin). Debris was cleared by centrifugation (20,000 × g, 30 min, 4°C). Streptomycin sulfate (1% wt/vol) was added dropwise over 20 min with gentle stirring, and precipitated material was cleared by centrifugation (20,000 × g, 30 min, 4°C). The lysate then was incubated with nickel resin (Qiagen) for 1 h with gentle agitation. The resin was washed progressively with wash buffers (20 mM Tris pH 7.5; 150 mM KCl; 2 mM MgCl_2_; 50, 100, 200 mM imidazole; and 5 mM βME). Protein was eluted with elution buffer [20 mM Tris pH 7.5, 150 mM KCl, 2 mM MgCl_2_, 300 mM Imidazole (Fisher), 5 mM βME], concentrated, and loaded onto an SEC column (ÄKTA purification system, GE Healthcare; Hiload 26/600 Superdex 75 prep grade) and run in storage buffer (20 mM Tris pH 7.5, 200 mM KCl, 2 mM MgCl_2_, 5% glycerol, 5 mM TCEP). The protein was collected, concentrated, aliquoted, flash frozen in liquid nitrogen, and stored at – 80°C in single-use aliquots to avoid freeze/thaw.

### Expression and purification of His_6_-AUF1

BL21-CodonPlus cells (Agilent) were transformed with pET28a His_6_-AUF1 (p37) and grown as described above. The purification procedure was the same as for His_6_HuR except for the buffers: lysis buffer [50 mM NaH2PO4 pH 7.6, 300 mM NaCl, 5 mM imidazole, 5 mM βME, and 1 mM PMSF, 0.5% Nonidet P40, 0.15 mg/ml lysozyme, 0.1 mg/ml RNase A (Sigma)]; wash buffers (50 mM NaH_2_PO_4_ pH 7.6; 150 mM NaCl; 20, 50, 100, mM imidazole; and 5 mM βME); elution buffer (50 mM NaH_2_PO_4_ pH 7.6, 150 mM NaCl, 250 mM Imidazole, 5 mM βME). The eluate was supplemented with 0.1 mg/ml RNase A and rotated end-over-end for 1 h at room temperature. The resulting mixture was then dialyzed against storage buffer (50 mM HEPES pH 7.6, 150 mM KCl, 2 mM MgCl_2_, 2 mM TCEP) for 2 h at 4°C, then dialysis buffer was changed and dialysis was allowed to proceed overnight at 4°C. The protein was then concentrated and stored (with 5% glycerol added to storage buffer) at –80°C in single-use aliquots to avoid freeze/thaw.

### ^32^P-End-labeling of RNA oligos

RNA oligos (IDT; Table S8) were resuspended in RNase-free water to 200 μM. RNA was labeled with γ-^32^P ATP (Perkin Elmer) in a reaction containing (final concentrations) 20 pmol RNA, 20 pmol γ-^32^P ATP, 1X polynucleotide kinase buffer (NEB), and 0.4 U/μl polynucleotide kinase (NEB) in a total volume of 50 μl. The reaction was incubated at 37°C for 1 h, after which 50 μl of RNase-free water and 300 μl of Trizol LS were added and the mixture was left at room temperature for 5 min. 200 μl of CHCl_3_ were added, the mixture was shaken for 15 s, left at room temperature for 2 min, and then spun down briefly to collect the contents. The aqueous layer was collected and mixed with an equal volume of CHCl_3_, shaken, and spun down briefly. The aqueous layer was again collected, supplemented with 15 μg of GlycoBlue (Ambion) and ammonium acetate to a final concentration of 500 mM, and 3 volumes of 100% EtOH were added. The mixture was vortexed and allowed to precipitate overnight at –20°C, then centrifuged (20,000 × g, 15 min, 4°C). The labeled RNA pellet was washed twice with 75% EtOH in RNase-free water, dried briefly, and resuspeded in RNase-free water. Labeled RNA was stored at –80°C in single-use aliquots to avoid freeze/thaw.

### Electrophoretic Mobility Shift Assays

These experiments were modeled on previously reported procedures ^69^. End-labeled RNA probes were mixed 1:10 with unlabeled probe. The binding reaction contained (final concentrations) 10 mM Tris pH 7.5, 100 mM KCl, 1 mM EDTA, 0.1 mM DTT, 5% glycerol, 0.01 mg/ml BSA, 0.5 nM RNA, and varying concentrations of HuR or AUF1^p37^. This mixture was incubated for 30 min at room temperature. The gel (10% 71:1 acrylamide:bisacrylamide TBE gel) was pre-run at 220V for 20 min in chilled 0.5X TBE buffer. The reactions were then loaded and run at 220 V for 12 min. The gels were exposed to an IP screen (GE Healthcare) at –80°C for 24–36 h, then imaged on a Typhoon FLA 7000 IP Imager (GE Healthcare). The bound RNA signal was quantitated using the “Gel Analysis” tool in ImageJ and fit to a one-site binding model (Equation 1 below) using Prism 7.

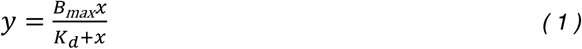

where y is the bound RNA signal, B_max_ is the plateau, x is the concentration of the mRBP, and *K*_d_ is the dissociation constant.

## Acknowledgments

American Heart Association predoctoral fellowship (17PRE33670395) (to J.R.P.); Swiss National Science Foundation (SNSF); Swiss Federal Institute of Technology Lausanne (EPFL); Novartis Medical-Biological Research Foundation (Switzerland); NCCR Chemical Biology (Switzerland); NIH Director’s New Innovator (1DP2GM114850) (to Y.A.). Michael T. Disare (a former undergraduate researcher in the Aye Lab) for assistance with cloning plasmids and protein purification; Dr. Xuyu Liu (a former postdoctoral researcher in the Aye Lab) for HNE(alkyne); Dr. Sung-Hee Chang and Prof. Timothy Hla (Weill Cornell Medicine and Harvard Medical School) for discussions of HuR biology.

## Author Contributions

J.R.P. Investigation, Formal analysis, Writing (initial draft and later editing), Visualization; M.J.C.L. Investigation (initial shRNA experiments, microinjection of zebrafish embryos), Formal analysis, Writing (critical review and editing); Y.A. Conceptualization, Writing, Supervision, Project administration, Funding acquisition. All authors assisted with final proofing of the manuscript.

## Note

RNA-Sequencing data have been submitted to the Gene Expression Omnibus (GEO; accession number GSE127444).

