## Supplementary material for "Post-transcriptional regulation of Nrf2-mRNA by the mRNA-binding proteins HuR and AUF1": SI

Jesse R. Poganik[§,†], Marcus J. C. Long[%], Michael T. Disare[†], Xuyu Liu[§], Sung-Hee Chang[#], Timothy Hla[‡,¶], and Yimon Aye[§,*]

[§] Swiss Federal Institute of Technology Lausanne (EPFL), 1015 Lausanne, Switzerland

[†]Department of Chemistry and Chemical Biology, Cornell University, Ithaca, NY 14853

[%]47 Pudding Gate, Bishop Burton, Beverley, HU17 8QH, UK

[#]Department of Medicine, Weill Cornell Medicine, New York, NY 10065

[‡]Department of Surgery, Harvard Medical School, Boston, MA 02115

[¶]Vascular Biology Program, Boston Children's Hospital, Boston, MA 02115

**RNA sequencing.** Following treatment of shHuR/shControl cells with HNE (50 μM) or $H_2O_2$ (225 μM) for 18 h, cells were lysed by the addition of Trizol (Invitrogen) and RNA was isolated following the manufacturer's protocol. The quality of the RNA was assessed by Nanodrop spectrophotometry (A260/A280 ratio ~2) and fragment analysis using a Bioanalyzer (Agilent). RNA quality numbers (RQNs) for all samples were 10.0. The rest of the procedure including data analysis was performed by the RNA sequencing core (RSC) service at Cornell University. Briefly, ribosomal (r)RNA was subtracted by hybridization from total RNA samples (100 ng total RNA input) using the RiboZero Magnetic Gold H/M/R Kit (Illumina). Following cleanup by precipitation, rRNA-subtracted samples was quantified with a Qubit 2.0 (RNA HS kit; Thermo Fisher). TruSeq-barcoded RNAseq libraries were generated from half of the rRNA-subtracted samples with the NEBNext Ultra II Directional RNA Library Prep Kit (New England Biolabs). Each library was quantified with a Qubit 2.0 (dsDNA HS kit; Thermo Fisher) and the size distribution was determined with a Fragment Analyzer (Advanced Analytical) prior to pooling. Libraries were sequenced on a NextSeq500 instrument (Illumina). At least 60M single-end 75bp reads were generated per library. Reads were trimmed for low quality and adaptor sequences with cutadapt v1.8 [parameters: –m 50 –q 20 –a AGATCGGAAGAGCACACGTCTGAACTCCAGTC –match-read-wildcards (Martin, 2011)]. Trimmed reads were mapped to rRNA sequences with bowtie2 v2.2 to remove them [Parameters: default mapping options, –al and –un to split matching and non-matching reads (Langmead and Salzberg, 2012)]. Reads were mapped to the reference genome/transcriptome (GRCh37/hg19]) using tophat v2.1 [parameters: –library-type=fr-firststrand –no-novel-juncs –G <ref_genes.gtf> (Kim et al., 2013)]. Cufflinks v2.2 (cuffnorm/cuffdiff) was used to generate FPKM values and statistical analysis of differential gene expression (Trapnell et al., 2013).

$$y = \frac{B_{max}x}{K_d + x} \qquad (1)$$

where y is the bound RNA signal, $B_{max}$ is the plateau, x is the concentration of the mRBP, and $K_d$ is the dissociation constant.

### Supplemental Tables

**Table S1. Comparison of HuR dissociation constants and HuR PAR-CLIP conversions for selected HuR targets.**

| **mRNA**[a] | | ***K*****$_d$** | **Total HuR PAR-CLIP nucleotide conversions** | **References** |
|---|---|---|---|---|
| COX2 | Site 1 | 20 nM | 59 | (Sengupta et al., 2003)<br>(Lebedeva et al., 2011) |
| | Site 2 | 18 nM | | |
| | Site 3 | 20 nM | | |
| c-Fos | (fragment) | 10 nM | 4 | |
| CXCL2 | (fragment) | 1.8 µM | 7 | (Ke et al., 2017)<br>(Lebedeva et al., 2011) |
| VEGFA | (fragment) | 30 nM | 113 | (Li et al., 2009)<br>(Lebedeva et al., 2011) |

[a] Selected mRNAs are those for which binding data is available in the literature.

**Table S2. Sequences of shRNAs targeting HuR and AUF1 and control hairpins.**

| Name | Target Gene | Sigma Serial Number | Sequence |
|---|---|---|---|
| shHuR (human) | ELAVL1 (human HuR) | TRCN0000017277 | 5′–CCGGCCCATCACAGTGAAGTTTGCACTCGAGTGCAAACTTCACTGTGATGGGTTTTT–3′ |
| shHuR (1) (mouse) | Elavl1 (mouse HuR) | TRCN0000112087 | 5′–CCGGCATTGGGAGAACGAATTTAATCTCGAGATTAAATTCGTTCTCCCAATGTTTTTG–3′ |
| shmHuR (2) (mouse) | Elavl1 (mouse HuR) | TRCN0000308990 | 5′–CCGGCGAGGTTGAATCTGCAAAGCTCTCGAGAGCTTTGCAGATTCAACCTCGTTTTTG–3′ |
| shAUF1 (1) (human) | HNRNPD(AUF1) | TRCN0000293283 | 5′–CCGGTCGAAGGAACAATATCAGCAACTCGAGTTGCTGATATTGTTCCTTCGATTTTTG–3′ |
| shAUF1 (2) (human) | HNRNPD(AUF1) | TRCN0000001293 | 5′–CCGGAGAGTGGTTATGGGAAGGTATCTCGAGATACCTTCCCATAACCACTCTTTTTT–3′ |
| shControl | GFP | N/A | 5′– GTCGAGCTGGACGGCGACGTA–3′ |

**Table S3. Raw counts, processed FPKM values, and Cuffdiff pairwise comparison test results from RNA-seq.**

(Please see excel file)

**Table S4. Descriptive statistics of RNA-seq fold changes for pairwise comparisons.**

| | shHuR (HNE treated/ untreated) ($Log_2$) | shControl (HNE treated/ untreated) ($Log_2$) | shHuR ($H_2O_2$ treated/ untreated) ($Log_2$) | shControl ($H_2O_2$ treated/ untreated) ($Log_2$) | shHuR/ shControl (no treatment) ($Log_2$) |
|---|---|---|---|---|---|
| **Minimum** | -5.0 | -2.9 | -3.6 | -3.9 | -3.9 |
| **25% Percentile** | -0.16 | -0.10 | -0.09 | -0.09 | -0.17 |
| **Median** | -0.0030 | 0.0067 | 0.0005 | 0.0031 | 0.0001 |
| **75% Percentile** | 0.14 | 0.11 | 0.08 | 0.08 | 0.17 |
| **Maximum** | 2.6 | 3.3 | 3.5 | 4.1 | 4.9 |
| **Range** | 7.6 | 6.2 | 7.0 | 8.0 | 8.8 |
| **Mean** | -0.034 | 0.010 | -0.017 | -0.012 | -0.008 |
| **Std. Deviation** | 0.36 | 0.25 | 0.26 | 0.32 | 0.36 |
| **Std. Error of Mean** | 0.0032 | 0.0022 | 0.0022 | 0.0026 | 0.0030 |
| **Skewness** | -2.9 | 1.8 | -0.7 | -0.8 | -0.3 |

Columns are color-coded to synchronize with the color scheme deployed in Figure S3.

**Table S5. Comparison of RNA-seq fold changes and HuR PAR-CLIP conversion events.**

| **Genes SDE in shHuR cells upon HNE treatment** | | | | | | | |
|---|---|---|---|---|---|---|---|
| **Gene** | **Fold change (HNE/non-treated)** | **q-value** | | | | | **Total PAR-CLIP conversions[a]** |
| ME1 | 1.6 | 0.04 | | | | | 28 |
| TXNRD1 | 1.6 | 0.02 | | | | | 1271 |
| FTL | 1.9 | 0.02 | | | | | ND |
| HMOX1 | 2.6 | 0.02 | | | | | 9 |
| **Genes SDE in shControl cells upon HNE treatment** | | | | | | | |
| **Gene** | **Fold change (HNE/non-treated)** | **q-value** | | | | | **Total PAR-CLIP conversions[a]** |
| PIR | 1.9 | 0.02 | | | | | 8 |
| **Genes SDE upon HuR knockdown alone** | | | | | | | |
| **Gene** | **Fold change (shHuR/shControl, non-treated)** | **q-value** | **Fold change (shHuR/shControl, HNE treatment)** | **q-value** | **Fold change (shHuR/shControl, $H_2O_2$ treatment)** | **q-value** | **Total PAR-CLIP conversions[a]** |
| SLC2A3 | 0.44 | 0.02 | 0.38 | 0.02 | 0.31 | 0.02 | 13 |
| MGST1 | 0.41 | 0.02 | 0.44 | 0.02 | 0.5 | 0.02 | 27 |
| INSIG1 | 0.54 | 0.02 | 0.54 | 0.02 | 0.57 | 0.02 | 125 |
| HSPA1B | 1.9 | 0.02 | 1.6 | 0.02 | 1.9 | 0.02 | ND |

[a](Lebedeva et al., 2011)
ND, not detected

**Table S6. Sequences of siRNAs targeting HuR and Nrf2.**

| Name | Target Gene | Dharmacon Serial Number | Sequence |
|---|---|---|---|
| siHuR (1) | ELAVL1 (HuR) | D-003773-04 | 5′–UCAAGACGCCAACUUGUA–3′ |
| siHuR (2) | ELAVL1 (HuR) | D-003773-05 | 5′–CAAAGACGCCAACUUGUAC–3′ |
| siNrf2 (1) | NFE2L2 (Nrf2) | D-003755-02 | 5′–CCAAAGAGCAGUUCAAUGA–3′ |
| siNrf2 (2) | NFE2L2 (Nrf2) | D-003755-04 | 5′–UAAAGUGGCUGCUCAGAAU–3′ |
| siNrf2 (3) | NFE2L2 (Nrf2) | D-003755-05 | 5′–UGACAGAAGUUGACAAUUA–3′ |

**Table S7. Sequences of morpholino (MO) oligonucleotides.**

| MO | Target Gene | Sequence |
|---|---|---|
| HuR ATG-MO | *elavl1* (HuR) | 5′–TGTGGTCTTCGTAACCGTTCGACAT–3′ |
| AUF1-ATG-MO | *hnrnpd* (AUF1) | 5'–ACCCAGAAACTGCTCCTCCGACATA–3' |
| AUF1-SPL-MO | *hnrnpd* (AUF1) | 5'–ACCTAGGATTATTTCGAGTCAGGAT–3' |
| Random Control MO | (non-targeting) | 5′–NNNNNNNNNNNNNNNNNNNNNNNNN–3′ |

**Table S8. Sequences of RNA oligos used for EMSA experiments.**

| | |
|---|---|
| Nrf2 3′–UTR Site 1 | 5′–GCUAGUUUUUUUGUACUAUU–3′ |
| Nrf2 3′–UTR Site 1 negative control | 5′–GCGAGCGCGCGCACGAGC–3′ |
| Nrf2 3′–UTR Site 2 | 5′–AAAAACUUAUUUAUACUGUU–3′ |
| Nrf2 3′–UTR Site 2 negative control | 5′– AAAAACGCACGCACACGGCG–3′ |
| Nrf2 3′–UTR Site 3 | 5′–AAAAAAAUUUUAAGAGCUGG–3′ |
| Nrf2 3′–UTR Site 3 negative control | 5′–AAAAAAAGCGCAAGAGCCGG–3′ |

**Table S9. Cloning primers.**

| Plasmid | Primer | Sequence |
|---|---|---|
| pCS2+8 Flag$_3$HuR | Fwd | 5′–TAATTAAAGGCCGGCCAGCGATCGC CGGACATGGATTATAAAGATCATGATGGCG–3′ |
| | FwdExt | 5′–GCTACTTGTTCTTTTTGCAGGATCCACTAGT GGCGCGCCATTAATTAAAGGCCGGCCAGC–3′ |
| | Rev | 5′–TTCTAGAGGCTCGAGAGGCCTTGAATTCGA TTATTTGTGGGACTTGTTGGTTTT–3′ |
| | RevExt | 5′–CTTATCATGTCTGGATCTACGTAATACGACTC ACTATAGTTCTAGAGGCTCGAGAGGCCT–3′ |
| pCS2+8 Flag$_2$AUF1$^{p42}$ | Fwd1 | 5′–ACGACGATAAGGATTACAAAGACGATGATGA CAAAGGGTCCATGTCGGAGGAGCAGTTCG–3′ |
| | Fwd2 | 5′–GCCGGACATCGATTCCCACCATGGACTATAA GGATGACGACGATAAGGATTACAAAGACG–3′ |
| | FwdExt | 5′–ATCCACTAGTGGCGCGCCATTAATTAAAGGCC GGCCAGCGATCGCCGGACATCGATTCCC–3′ |
| | Rev1 | 5′–CACTATAGTTCTAGAGGCTCGAGAGGCCTTT TAGTATGGTTTGTAGCTATTTTGATGACC–3′ |
| | Rev2 | 5′–CACTATAGTTCTAGAGGCTCGAGAGG–3′ |
| | RevExt | 5′–CTTATCATGTCTGGATCTACGTAATACGACTC ACTATAGTTCTAGAGGCTCGAGAGGCCT–3′ |
| pCS2+8 Flag$_2$AUF1$^{p40/p45}$ | Fwd1 | 5′–CTCTGAAGCAGCGACGGCACAGCGGGAAGAAT GGAAAATGTTTATAGGAGGCCTTAGCTG–3′ |
| | Fwd2 | 5′–GAACGAGGAGGATGAAGGCCATTCAAACTCCT CCCCACGACACTCTGAAGCAGCGACGGC–3′ |
| | FwdExt | 5′–GGCAGCGCCGAGTCGGAGGGGGCGAAGATTG ACGCCAGTAAGAACGAGGAGGATGAAGGC–3′ |
| | Rev | 5′–CTTCGACATGGCTACTTTTATTTCA–3′ |
| pCS2+8 Flag$_2$AUF1$^{p37}$ | Fwd | 5′–ATTTAGGTGACACTATAG–3′ |
| | Rev | 5′–TGGATACCTTCCCATAACCACTCTGCTGGTCAC CACCTCTTCCACGAGCTCTTCCTGCAA–3′ |
| | RevExt | 5′–TAGTATGGTTTGTAGCTATTTTGATGACC ACCTCGCCTGGATACCTTCCCATAACCACTC–3′ |
| pSGG-Nrf2 intron 2 reporter | Fwd | 5′–CAATCCGGTACTGTTGGTAAAGCCACCATG GACATGGATTTGATTGACATACTTTG–3′ |
| | FwdExt | 5′–TTTAGGTCTGTTCTCGTCTTCCGAGATCTAAGC TTGGCAATCCGGTACTGTTGGTAAAGC–3′ |
| | Rev | 5′–GCCCTTCTTAATGTTTTTGGCATCTTCCATCT CATTGTCATCTACAAACGGG–3′ |
| | RevExt | 5′–GGCGGTCCCGTCTTCGAGTGGGTAGAATGGC GCTGGGCCCTTCTTAATGTTTTTGGCATC–3′ |
| pSGG-Nrf2 intron 3 reporter | Fwd | 5′–CAATCCGGTACTGTTGGTAAAGCCACCATG GTTGCCCACATTCCCAAATC–3′ |
| | FwdExt | 5′–TTTAGGTCTGTTCTCGTCTTCCGAGATCTAAGC TTGGCAATCCGGTACTGTTGGTAAAGC–3′ |
| | Rev | 5′–GCCCTTCTTAATGTTTTTGGCATCTTCCATCTGTAACTCAGG AATGGATAATAGCTCC–3′ |

| | | |
|---|---|---|
| | RevExt | 5′–GGCGGTCCCGTCTTCGAGTGGGTAGAATGGC GCTGGGCCCTTCTTAATGTTTTTGGCATC–3′ |
| pSGG-Nrf2 intron 4 reporter | Fwd | 5′–CAATCCGGTACTGTTGGTAAAGCCACCATG GTTTCTTCGGCTACGTTTCAGTC–3′ |
| | FwdExt | 5′–TTTAGGTCTGTTCTCGTCTTCCGAGATCTAAGC TTGGCAATCCGGTACTGTTGGTAAAGC–3′ |
| | Rev | 5′–GCCCTTCTTAATGTTTTTGGCATCTTCCATCTGGT TGGGGTCTTCTGTGG–3′ |
| | RevExt | 5′–GGCGGTCCCGTCTTCGAGTGGGTAGAATGGC GCTGGGCCCTTCTTAATGTTTTTGGCATC–3′ |
| pET28a $His_6$AUF1 | Fwd | 5′–TGGTGCCTCGTGGTAGCCATATGGACTATAA GGATGACGACGATAAG–3′ |
| | FwdExt | 5′–ATGGGCAGCAGCCATCATCATCATCATCAC AGCAGCGGCCTGGTGCCTCGTGGTAGCCAT–3′ |
| | Rev | 5′–TTAGTATGGTTTGTAGCTATTTTGATGACCTAA CAAAGCCCGAAAGGAAGCTGAG–3′ |
| | RevExt | 5′–TATGCTAGTTATTCAGCGGTGGCAGCAGCCA ACTCAGCATCCTTTCGGGCTTTGTTA–3′ |
| pET28a $His_6$HuR | Fwd | 5′–GCAGCGGCGAAAACTTGTATTTCCAGGGCTC AGGGATGTCTAATGGTTATGAAGACCACA–3′ |
| | FwdExt | 5′–AGATATACCATGGGCAGCAGCCATCATCATCA TCATCACAGCAGCGGCGAAAACTTGTAT–3′ |
| | Rev | 5′–CTCAGCTTCCTTTCGGGCTTTGTTATTATTTG TGGGACTTGTTGGTTTT–3′ |
| | RevExt | 5′–TATGCTAGTTATTCAGCGGTGGCAGCAGCCA ACTCAGCATCCTTTCGGGCTTTGTTA–3′ |
| shRNA-resistant HuR mutagenesis | Fwd | 5′–AAACCCCCAGGTTCCTCTGAACCTATTACCGT CAAATTCGCAGCCAACCCCAACCAGAAC–3′ |
| | Rev | 5′–GTTCTGGTTGGGGTTGGCTGCGAATTTGACGGTAA TAGGTTCAGAGGAACCTGGGGGTTT–3′ |

**Table S10. Antibodies.**

| Antibody | Source | Dilution(s)[a] |
|---|---|---|
| Mouse monoclonal anti-HuR | Santa Cruz Biotechnology sc-5261 | 1:1000 (WB)<br>1:200 (IF, zebrafish) |
| Rabbit polyclonal anti-AUF1 | EMD Millipore 07-260 | 1:500 (WB, cells)<br>1:250 (WB, zebrafish) |
| Mouse monoclonal anti-FLAG M2 | Sigma F1804 | 1:2000 (WB) |
| Goat polyclonal anti-GFP (FITC-conjugated) | Abcam ab6662 | 1:500 (IF, zebrafish) |
| Mouse monoclonal anti-β-actin (peroxidase-conjugated) | Sigma A3854 | 1:40000 (WB, cells)<br>1:3000 (WB, zebrafish) |
| Rabbit polyclonal anti-goat IgG (Alexa Fluor 568-conjugated) | Abcam ab175707 | 1:1000 (IF secondary, zebrafish) |
| Donkey polyclonal anti-mouse IgG (Alexa Fluor 568-conjugated) | Abcam ab175472 | 1:1000 (IF secondary, zebrafish) |
| Goat polyclonal anti-rabbit IgG (HRP-conjugated) | Cell Signaling Technology #7074 | 1:3000 (WB secondary) |
| Goat polyclonal anti-mouse IgG (HRP-conjugated) | Abcam ab6789 | 1:8000 (WB secondary) |
| Anti-mouse IgG VeriBlot for IP secondary antibody (HRP-conjugated) | Abcam ab131368 | 1:1000 (IP WB secondary) |

[a]WB, western blot; IF, immunofluorescence; IP, immunoprecipitation

**Table S11. qRT-PCR primers.**

| Gene | Species | Primer Sequences |
|---|---|---|
| NFE2L2 (Nrf2) | Human | Forward 5′–TATCCATTCCTGAGTTACAGTGTC–3′<br>Reverse 5′–CTGTCAGTTTGGCTTCTGGAC–3′ |
| VEGFA | Human | Forward 5′–ATCTGCATGGTGATGTTGGA–3′<br>Reverse 5′–GGGCAGAATCATCACGAAGT–3′ |
| GAPDH | Human | Forward 5′–TGGAAGGACTCATGACCACAG–3′<br>Reverse 5′–CAGCTCAGGGATGACCTTGC–3′ |
| ACTB | Human | Forward 5′–ATCATTGCTCCTCCTGAGC–3′<br>Reverse 5′–CTGCTTGCTGATCCACATC–3′ |
| ZNF200 | Human | Forward 5′–GGCGCACGCTGAGCTTTAC–3′<br>Reverse 5′–GGCGCACGCTGAGCTTTAC–3′ |
| Firefly Luc Reporter | (Ectopic reporter) | Forward 5′–ACGCACATATCGAGGTGGAC–3′<br>Reverse 5′–CCAACACGGGCATGAAGAAC–3′ |
| *Renilla* Luc Reporter | (Ectopic reporter) | Forward 5′–AGCGGGAATGGCTCATATCG–3′<br>Reverse 5′–CACGTCCACGACACTCTCAG–3′ |

### Supplemental Figure Legends

**Figure S1. HuR and AUF1 levels are efficiently knocked down in HEK293T cells (A–C) and larval zebrafish (D–E), related to Figures 1–4.** (**A**, **B,** and **C**) Representative western blots showing knockdown level of HuR with shRNA, AUF1 with shRNA, and HuR with siRNA, respectively. *Inset below each:* Quantitation (mean±SEM of n=3 independent replicates for each set) of western blot signal measured using the Gel Analysis function of ImageJ. (**D**) Fertilized eggs were injected with Control- or HuR-ATG-MO at the single-cell stage, grown for 24 h, then fixed and stained with an antibody that detects zHuR and other zebrafish Hu family proteins. Scale bar, 500 µm. *Inset below:* Quantitation (mean ± SEM of n=5 fish per condition) of mean fluorescence intensity measured with the Measure tool of ImageJ. *Note that this represents only an estimate of HuR knockdown efficiency as the antibody also detects other Hu-family proteins*. (**E**) Total zebrafish lysate from fish injected with MOs targeting zAUF1 (of which there are only two isoforms in zebrafish) was subjected to western blot analysis. All p-values were calculated using Student's t-test.

**Figure S2. Growth inhibition of shHuR and shControl cells by HNE and $H_2O_2$, related to Figure 1.** shHuR and shControl HEK293T cells were treated with the indicated concentration of HNE (**A**) or $H_2O_2$ (**B**) for 48 h, after which viability was assessed with alamarBlue. Data are presented as mean ± SEM (n≥8 per point) and were fit with Prism. $EC_{50}$ for growth inhibition is presented with standard error.

**Figure S3. Distribution of RNA-seq data, related to Figure 1.** Histograms of binned values of $\log_2$(fold change) for expressed genes under the indicated conditions (Figure 1) fit with a Gaussian function with Prism, displaying (1) the greater spread observed under HNE-stimulation conditions compared to $H_2O_2$, and (ii) positive vs. negative skewness in the individual data sets. The curves are offset in the plot for clarity. The Gaussian function used was $y=A*e^{(-0.5*((x-m)/SD)^2)}$, where y is the number of values in the bin, A is the amplitude, x is the distance from the bin center, m is the mean, and SD is the standard deviation.

**Figure S4. Expression level of ectopic HuR in shHuR cells is near-endogenous levels, related to Figure 1.** HEK293T cells were transfected with plasmid encoding flag-HuR. Western blot was probed with anti-HuR antibody. *Inset*: Quantitation of HuR levels derived from the Gel Analysis tool of ImageJ.

**Figure S5. Mouse embryonic endothelial cells (MEECs) depleted of HuR similarly feature suppressed Nrf2 activity in the non-stimulated state and greater Nrf2 activity upregulation in the HNE-stimulated state, related to Figure 1.** (**A**) Western blot assessing knockdown levels of HuR in MEEC shHuR/shControl lines. *Inset at right*: Quantitation (mean±SEM, n=4 replicates) of western blot data derived from the Gel Analysis tool of ImageJ. (**B** and **C**) Nrf2 activity in non-stimulated (**B**) and HNE-treated [25 µM, 18 h (**C**)] MEEC lines (mean±SEM, n≥31 for **B** and n=8 for **C**). All p values calculated with Student's t-test.

**Figure S6. Nrf2 activity levels are proportional to Nrf2 protein levels; suppression of Nrf2 activity upon HuR-knockdown in non-stimulated cells and greater Nrf2 activity upregulation upon HNE stimulation is Nrf2-dependent, related to Figure 2.** (**A**) HEK293T were co-transfected with Nrf2 activity reporters (Figure 1E) and siNrf2 and Nrf2 activity was measured (mean±SEM of n≥15 independent replicates per condition). (**B**) HEK293T were co-transfected with Nrf2 activity reporters (Figure 1E) and siHuR, siNrf2, both, or siControl and Nrf2 activity was measured (mean±SEM of n=16 independent replicates per condition). (**C**) Similar experimental setup as (**A**), but cells were treated with HNE(alkyne) (50 µM, 18 h), after which Nrf2 activity was measured (mean±SEM of n=8 independent replicates per condition). All p values calculated with Student's t-test.

**Figure S7. HuR and AUF1 act independently on Nrf2-mRNA, related to Figures 2 and 3.** (**A**) Data from Figure 2C plotted to show the level of Nrf2 activity suppression upon HuR-knockdown with or without the simultaneous AUF1-knockdown. Data points were divided by the average Nrf2 activity in the corresponding siControl condition shown in Figure 2C (mean±SEM of n=4 independent replicates). (**B**) Same analysis as in **A** of data from Figure 3F (mean±SEM of n=4 independent replicates). All p values calculated with Student's t-test.

**Figure S8. An MO targeting the zebrafish Nrf2 homolog, nfe2l2a, reduces GFP expression in *Tg(gstp1:GFP)* reporter zebrafish, related to Figure 2.** Fertilized eggs were injected at the single-cell stage with Nrf2a-ATG-MO, which depletes the key zebrafish Nrf2 homolog, nfe2l2a (Nrf2a). Fish were then immunostained as described above and IF data was quantitated (mean±SEM of n=38 fish per condition) using the Measure tool of ImageJ. Scale bar, 500 µm. p value calculated with Student's t-test.

**Figure S9. HuR and AUF1 bind to sites in the 3′–UTR of Nrf2-mRNA with nanomolar affinity, related to Figure 3.**

(**A**) Schematic of the 3′–UTR of Nrf2-mRNA showing relative positions of the three identified HuR- and AUF1-binding sites. (**B** and **C**) Recombinant $His_6$-HuR (**B**) or $His_6$-AUF1$^{p37}$ (**C**) were incubated with the indicated $^{32}P$-end-labeled RNA oligo for 30 min at room temperature, then run on a polyacrylamide gel to resolve bound and unbound RNA. Gels were visualized using autoradiography. Dissociation constants were derived by quantitating the bound RNA signal (as indicated) using the Gel Analysis tool of ImageJ (mean±SEM of n=2 independent experiments for each RNA/protein pair) and fitting to a one-site binding model ($y=B_{max}*x/(K_d+x)$, where y is the bound mRNA signal, $B_{max}$ is the plateau, x is the concentration of mRBP, and $K_d$ is the dissociation constant) in Prism. HuR binding to Site 2 was not quantitated due to negligible binding. Note that although AUF1$^{p37}$ was chosen as a representative isoform, all isoforms bind to Nrf2-mRNA in cells to a similar degree (see Figure 3B).

**Figure S10. HuR knockdown reduces total Nrf2-mRNA levels, but does not significantly perturb its half-life; AUF1 knockdown does significantly perturb Nrf2-mRNA stability, related to Figure 3.** (**A**) qRT-PCR was used to assess the levels of endogenous Nrf2-mRNA in HEK293T cells transfected with siHuR/siControl (mean±SEM of n=4, 8, and 8 independent replicates for siHuR (1), siHuR (2), and siControl, respectively). (**B**) 3'–UTR reporter mRNA levels (firefly luciferase mRNA normalized to *Renilla* luciferase mRNA) were measured with qRT-PCR [mean±SEM, n=8 for siHuR (1) and siControl, n=7 for siHuR (2)]. (**C**) shHuR/shControl HEK293T were treated with actinomycin D for the indicated time, after which RNA was isolated and qRT-PCR was used to measure the amount of endogenous Nrf2-mRNA remaining. (**D**) shAUF1/shControl cells were treated as in **C** and qRT-PCR was used to measure the amount of endogenous Nrf2-mRNA remaining over time. *Inset at right*: Quantification (mean±SEM, n=4 for shAUF1 and n=3 for shControl) of Nrf2-mRNA half-life in shAUF1 and shControl cells. For **C** and **D**, data were fit to a one-phase decay model ($y = (y_0 - P)*e^{(-k*x)} + P$, where y is the amount of Nrf2-mRNA remaining, $y_0$ is the initial amount of Nrf2-mRNA, P is the plateau, $k$ is the rate constant, and x is the time) using Prism. Decay rates constants ($k$) are presented as mean±SD. All p values calculated with Student's t-test.

**Figure S11. Illustrations of premature non-functional vs. mature Nrf2-mRNA and detailed view of the former, related to Figure 4.** (**A**) Premature Nrf2-mRNA is not functional because it does not lead to production of Nrf2-protein. The premature mRNA must be spliced and exported from the nucleus to be translated to protein. (**B**) Schematic of premature Nrf2-mRNA. Boxes represent exons and lines represent introns. The size of each exon/intron in base pairs (bp) is shown for each (Figure 4).

### Figure S1

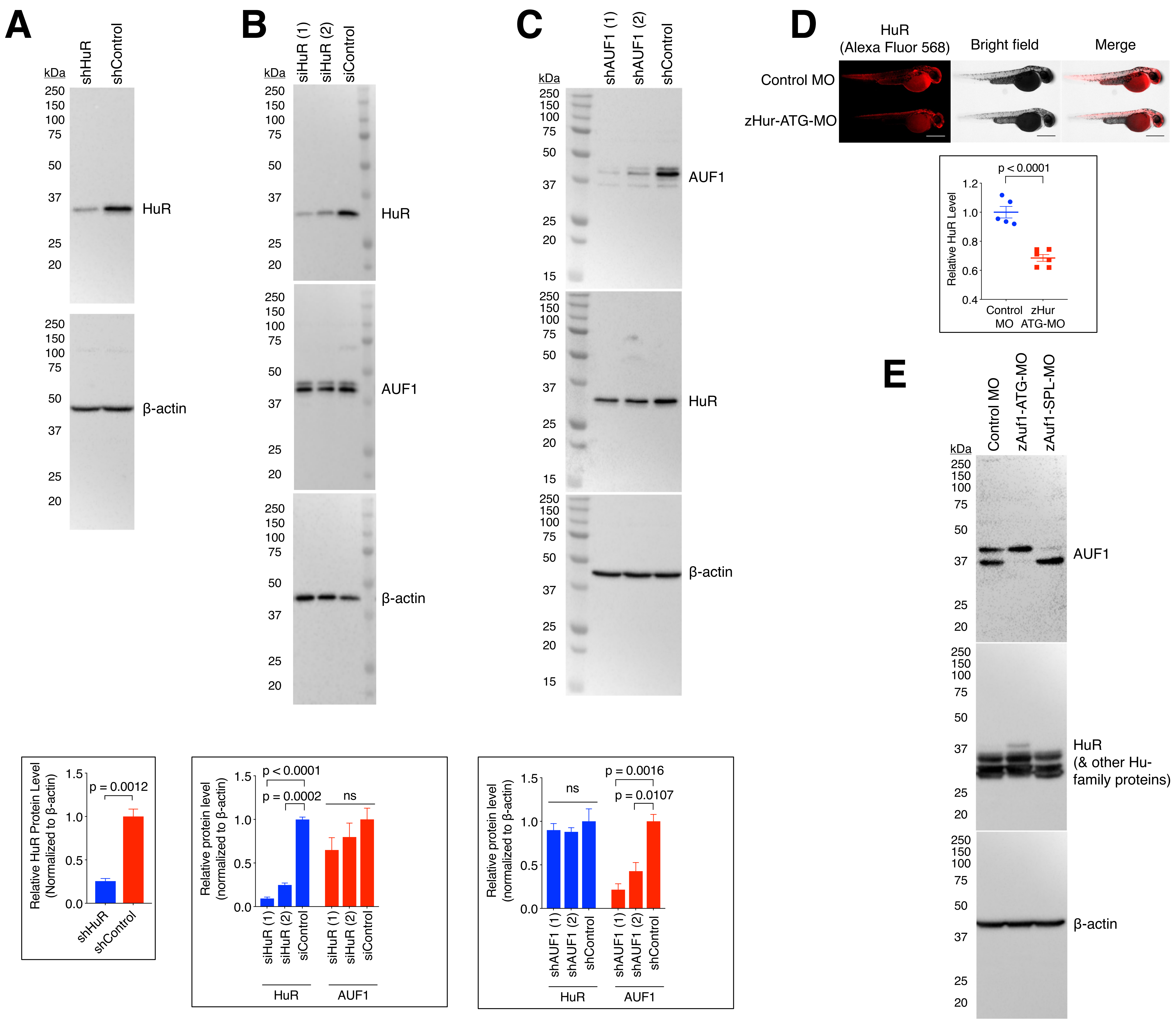

A
shHuR
shControl
kDa
HuR
β-actin
Relative HuR Protein Level (Normalized to β-actin)
p = 0.0012
B
siHuR (1)
siHuR (2)
siControl
HuR
AUF1
β-actin
Relative protein level (normalized to β-actin)
p < 0.0001
p = 0.0002
ns
C
shAUF1 (1)
shAUF1 (2)
shControl
AUF1
HuR
β-actin
p = 0.0016
p = 0.0107
ns
D
HuR (Alexa Fluor 568)
Bright field
Merge
Control MO
zHur-ATG-MO
Relative HuR Level
p < 0.0001
Control MO
zHur ATG-MO
E
Control MO
zAuf1-ATG-MO
zAuf1-SPL-MO
AUF1
HuR (& other Hu-family proteins)
β-actin

### Figure S2

A

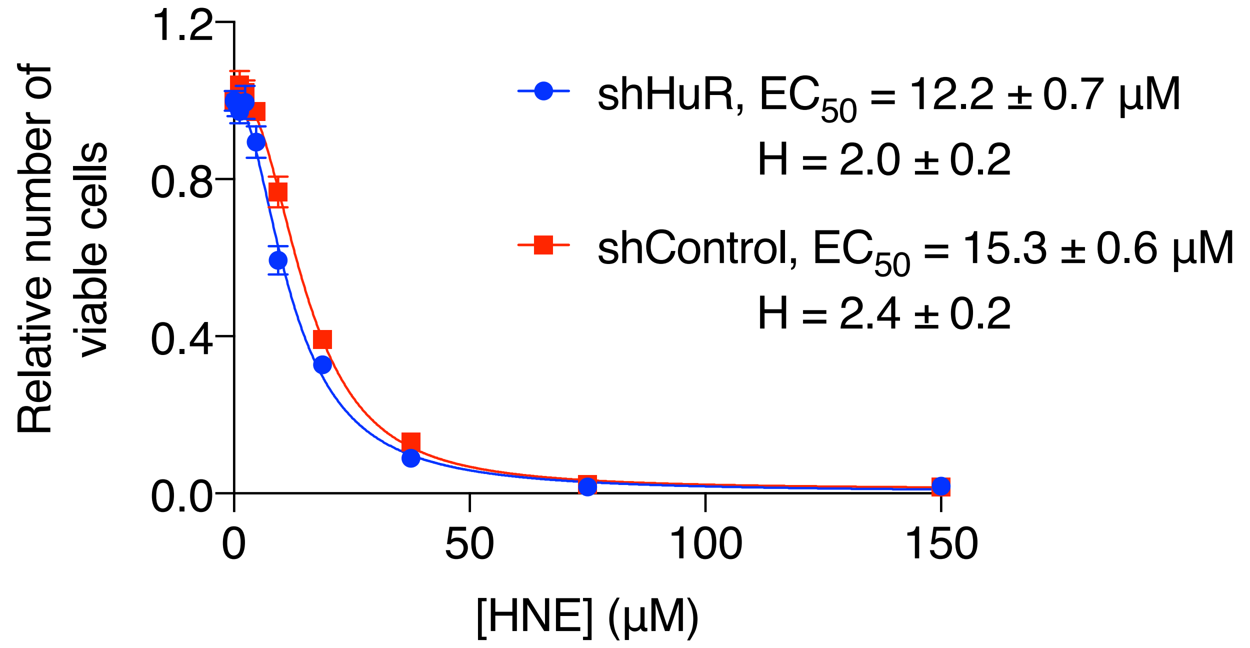

shHuR, EC50 = 12.2 ± 0.7 µM
H = 2.0 ± 0.2
shControl, EC50 = 15.3 ± 0.6 µM
H = 2.4 ± 0.2
Relative number of viable cells
[HNE] (µM)


B

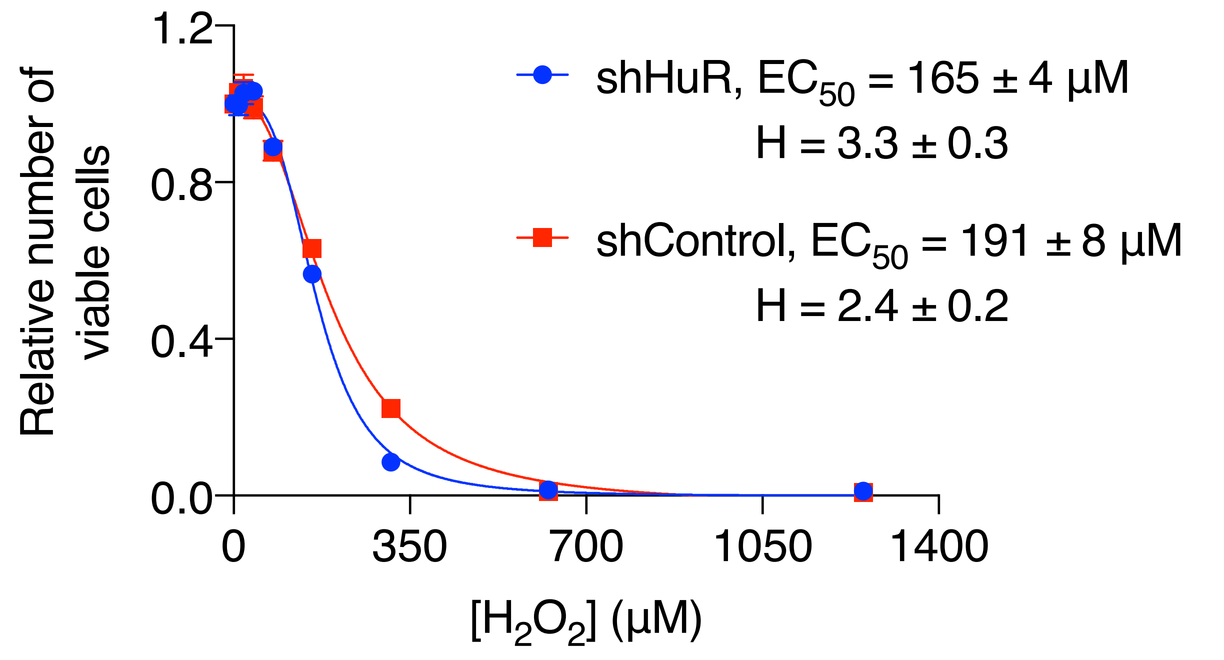

shHuR, EC50 = 165 ± 4 µM
H = 3.3 ± 0.3
shControl, EC50 = 191 ± 8 µM
H = 2.4 ± 0.2
Relative number of viable cells
[H2O2] (µM)

### Figure S3

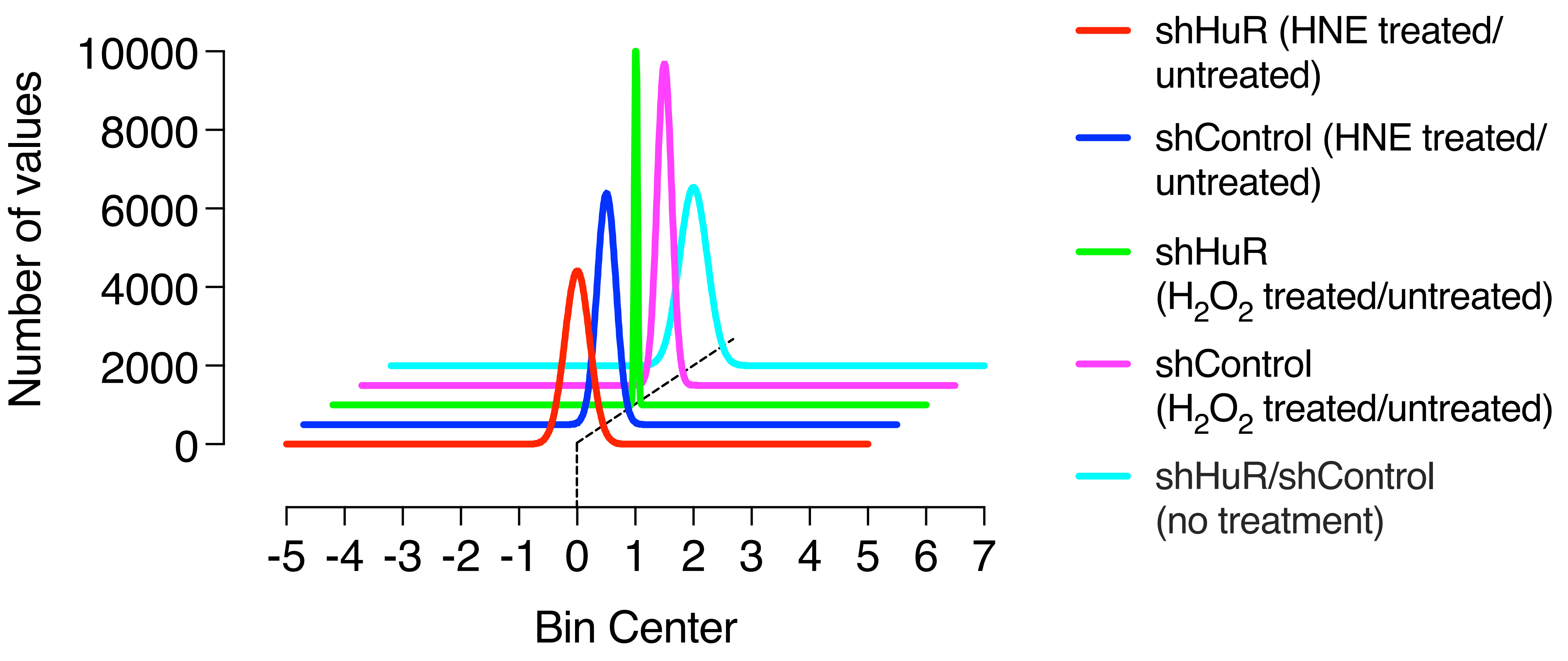

Number of values
10000
8000
6000
4000
2000
0
-5 -4 -3 -2 -1 0 1 2 3 4 5 6 7
Bin Center
shHuR (HNE treated/untreated)
shControl (HNE treated/untreated)
shHuR ($H_2O_2$ treated/untreated)
shControl ($H_2O_2$ treated/untreated)
shHuR/shControl (no treatment)

### Figure S4

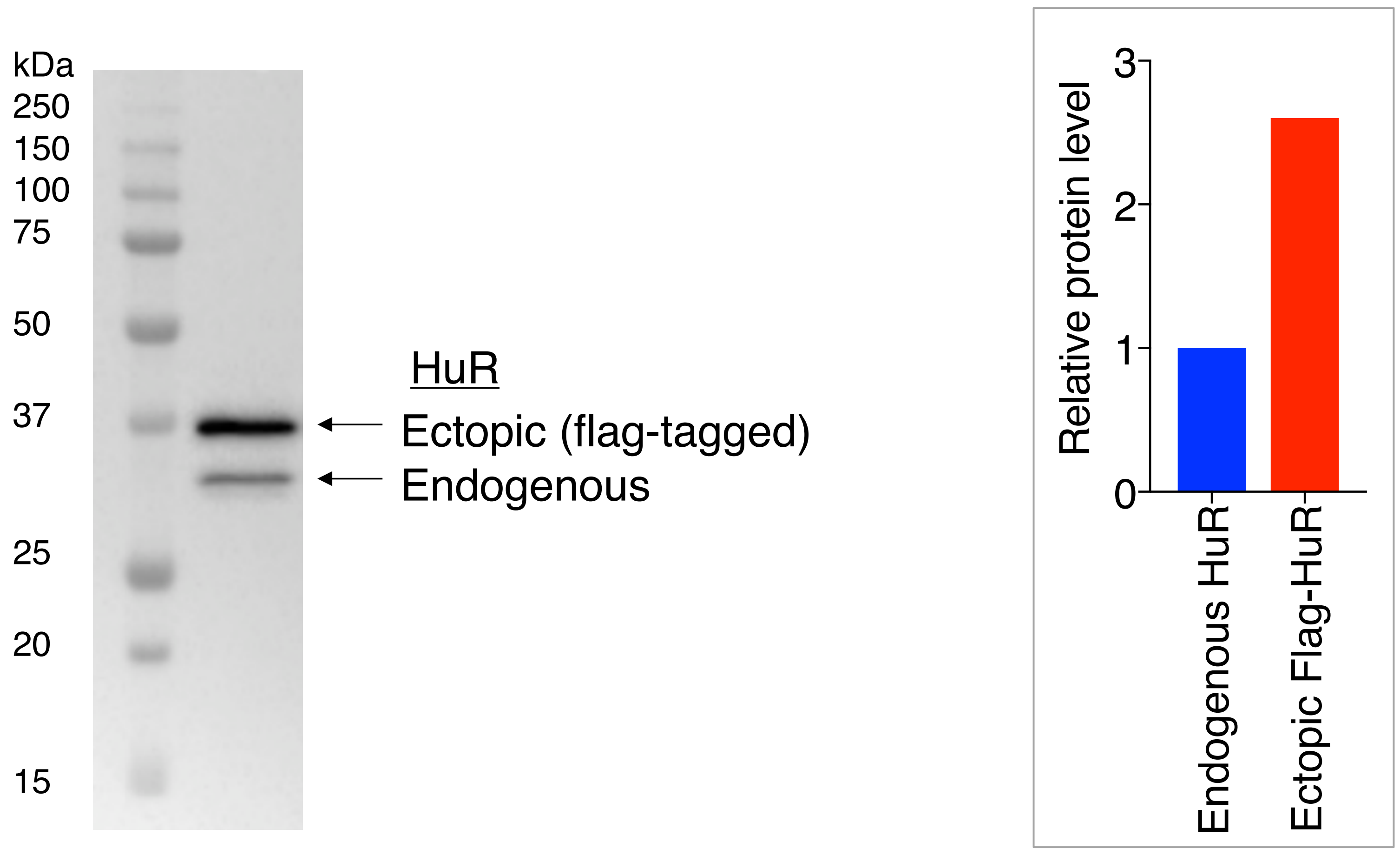

kDa
250
150
100
75
50
37
25
20
15
HuR
Ectopic (flag-tagged)
Endogenous
Relative protein level
3
2
1
0
Endogenous HuR
Ectopic Flag-HuR

### Figure S5

A

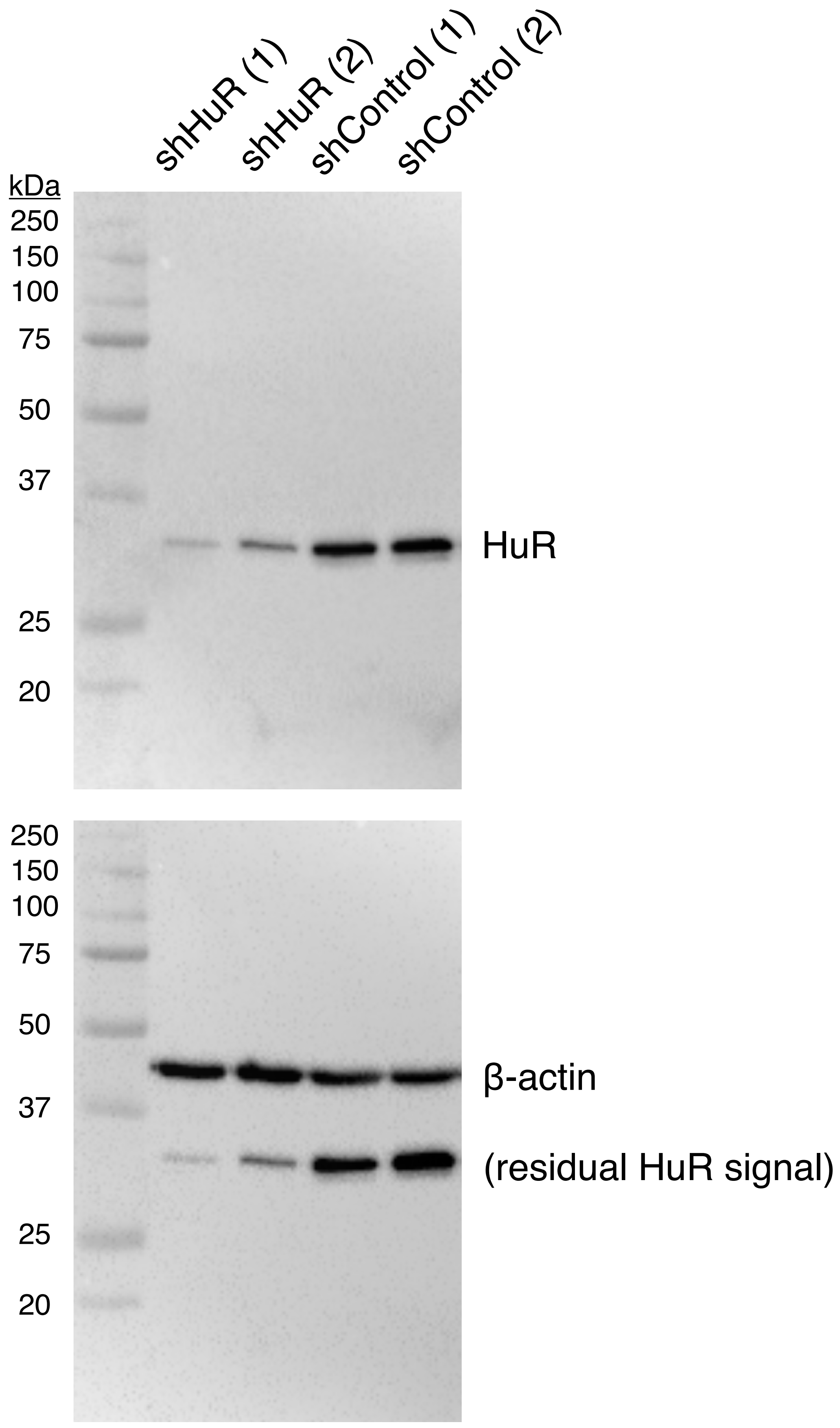


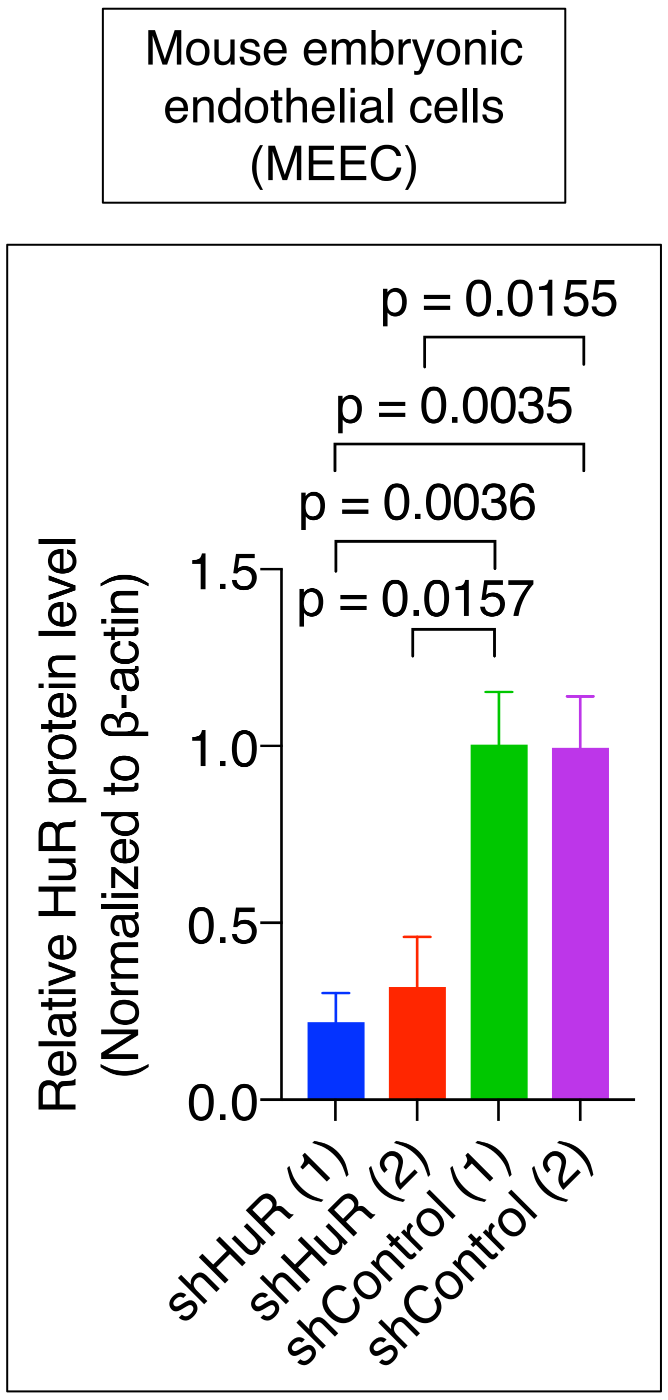


B

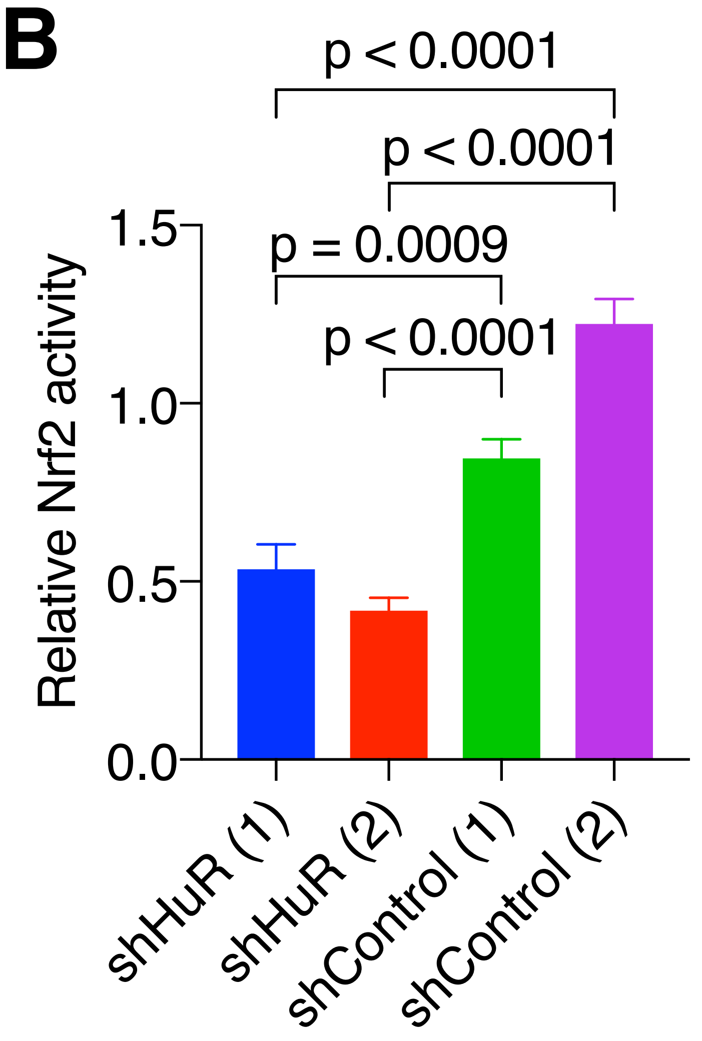


C

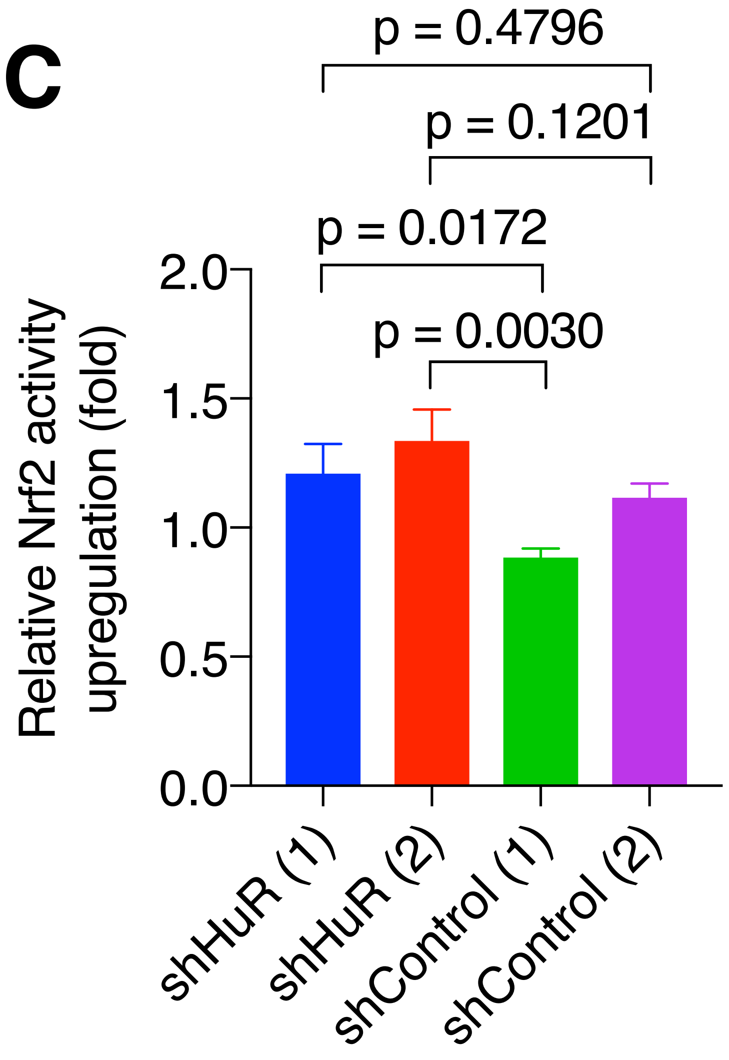

### Figure S6

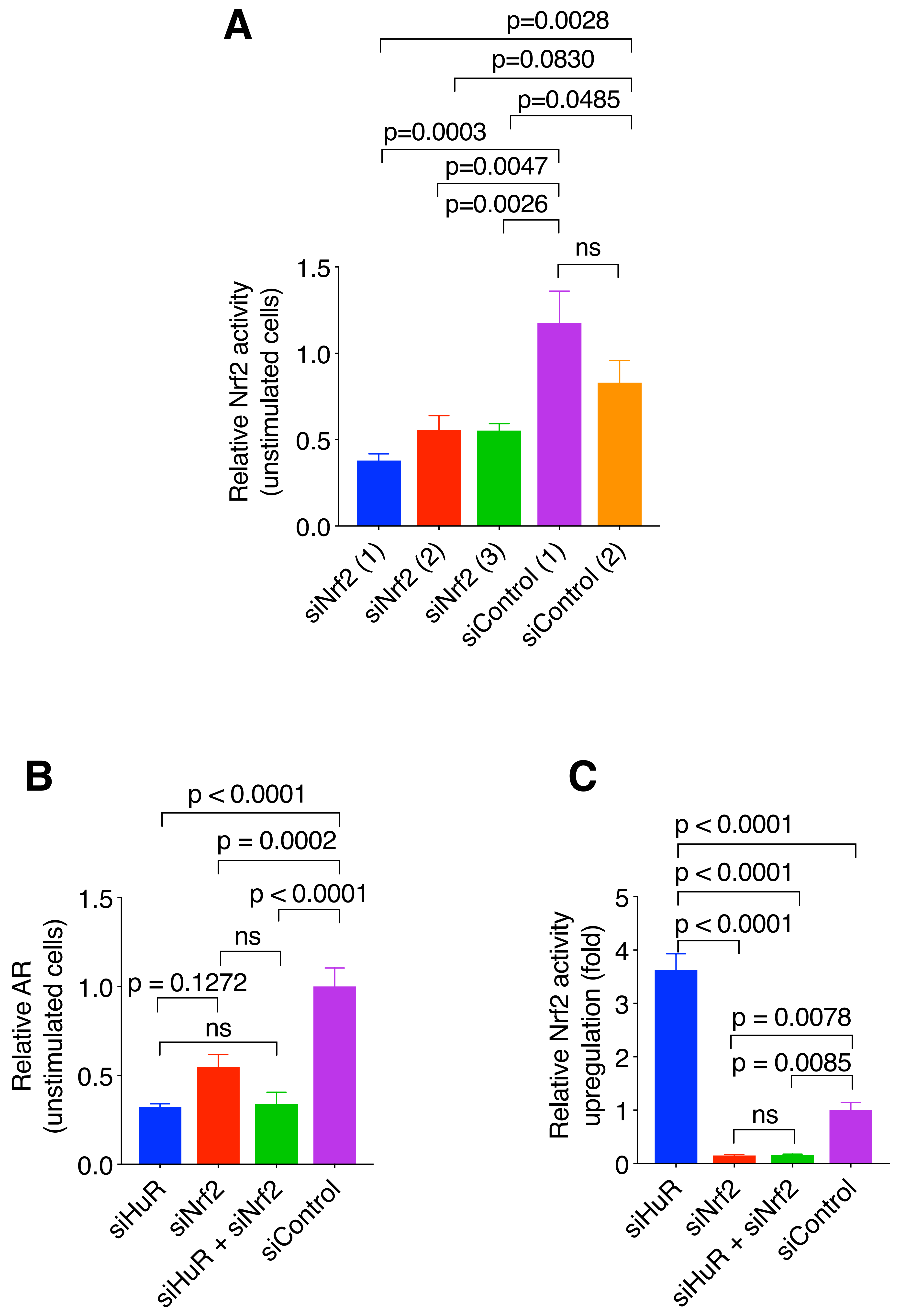
A
p=0.0028
p=0.0830
p=0.0485
p=0.0003
p=0.0047
p=0.0026
ns
Relative Nrf2 activity (unstimulated cells)
1.5
1.0
0.5
0.0
siNrf2 (1)
siNrf2 (2)
siNrf2 (3)
siControl (1)
siControl (2)
B
p < 0.0001
p = 0.0002
p < 0.0001
ns
p = 0.1272
ns
Relative AR (unstimulated cells)
siHuR
siNrf2
siHuR + siNrf2
siControl
C
p < 0.0001
p < 0.0001
p < 0.0001
p = 0.0078
p = 0.0085
ns
Relative Nrf2 activity upregulation (fold)
5
4
3
2
1
0
siHuR
siNrf2
siHuR + siNrf2
siControl

### Figure S7

A

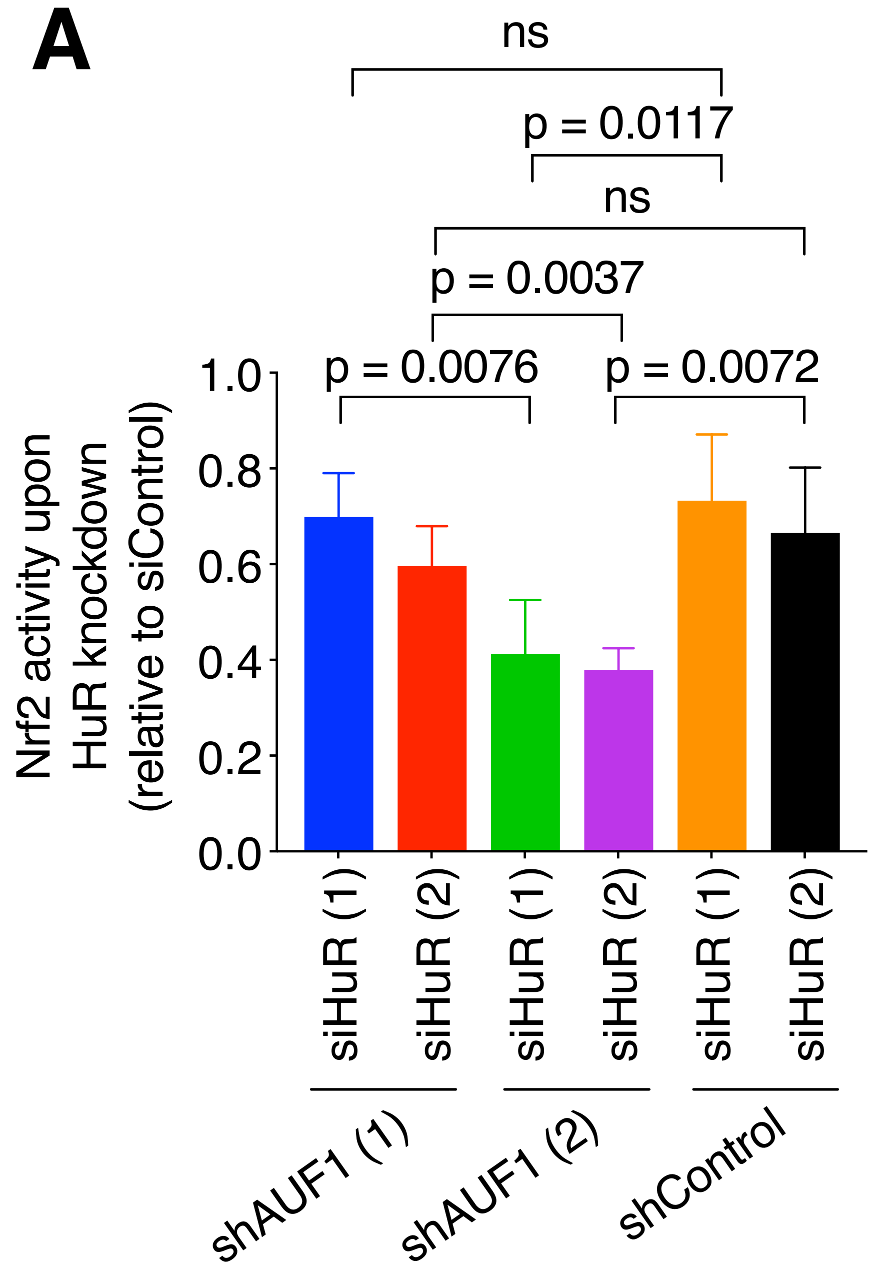
ns
p = 0.0117
ns
p = 0.0037
p = 0.0076
p = 0.0072
Nrf2 activity upon HuR knockdown (relative to siControl)
1.0
0.8
0.6
0.4
0.2
0.0
siHuR (1)
siHuR (2)
siHuR (1)
siHuR (2)
siHuR (1)
siHuR (2)
shAUF1 (1)
shAUF1 (2)
shControl

B

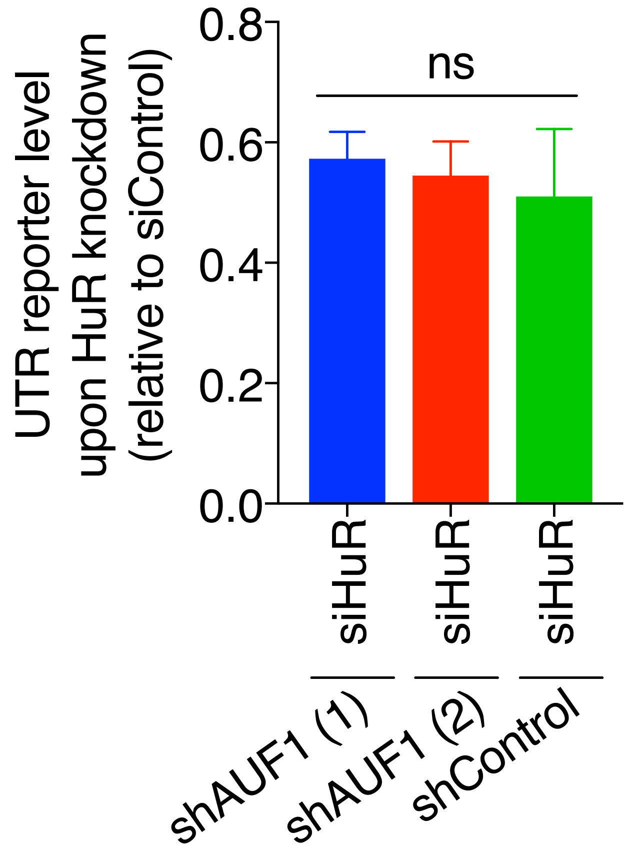
ns
UTR reporter level upon HuR knockdown (relative to siControl)
0.8
0.6
0.4
0.2
0.0
siHuR
siHuR
siHuR
shAUF1 (1)
shAUF1 (2)
shControl

### Figure S8

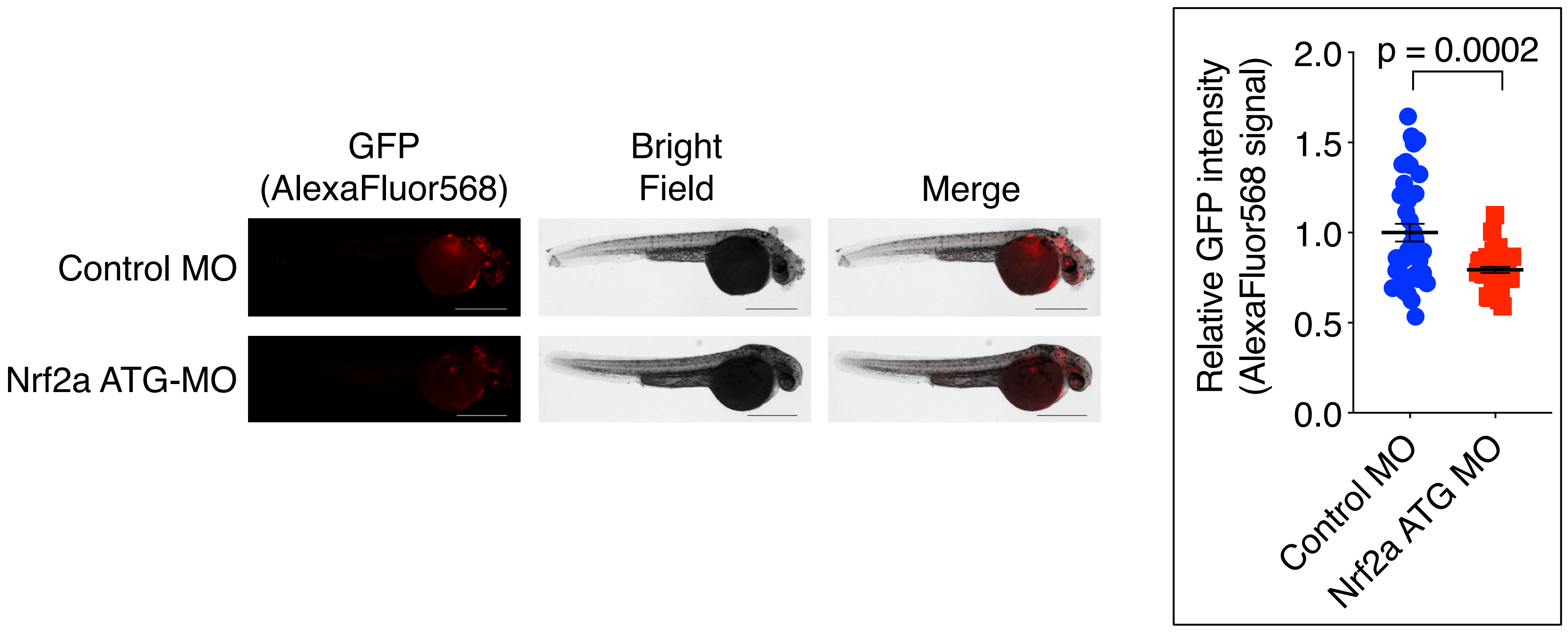
GFP
(AlexaFluor568)
Bright
Field
Merge
Control MO
Nrf2a ATG-MO
Relative GFP intensity
(AlexaFluor568 signal)
2.0
1.5
1.0
0.5
0.0
p = 0.0002
Control MO
Nrf2a ATG MO

### Figure S9

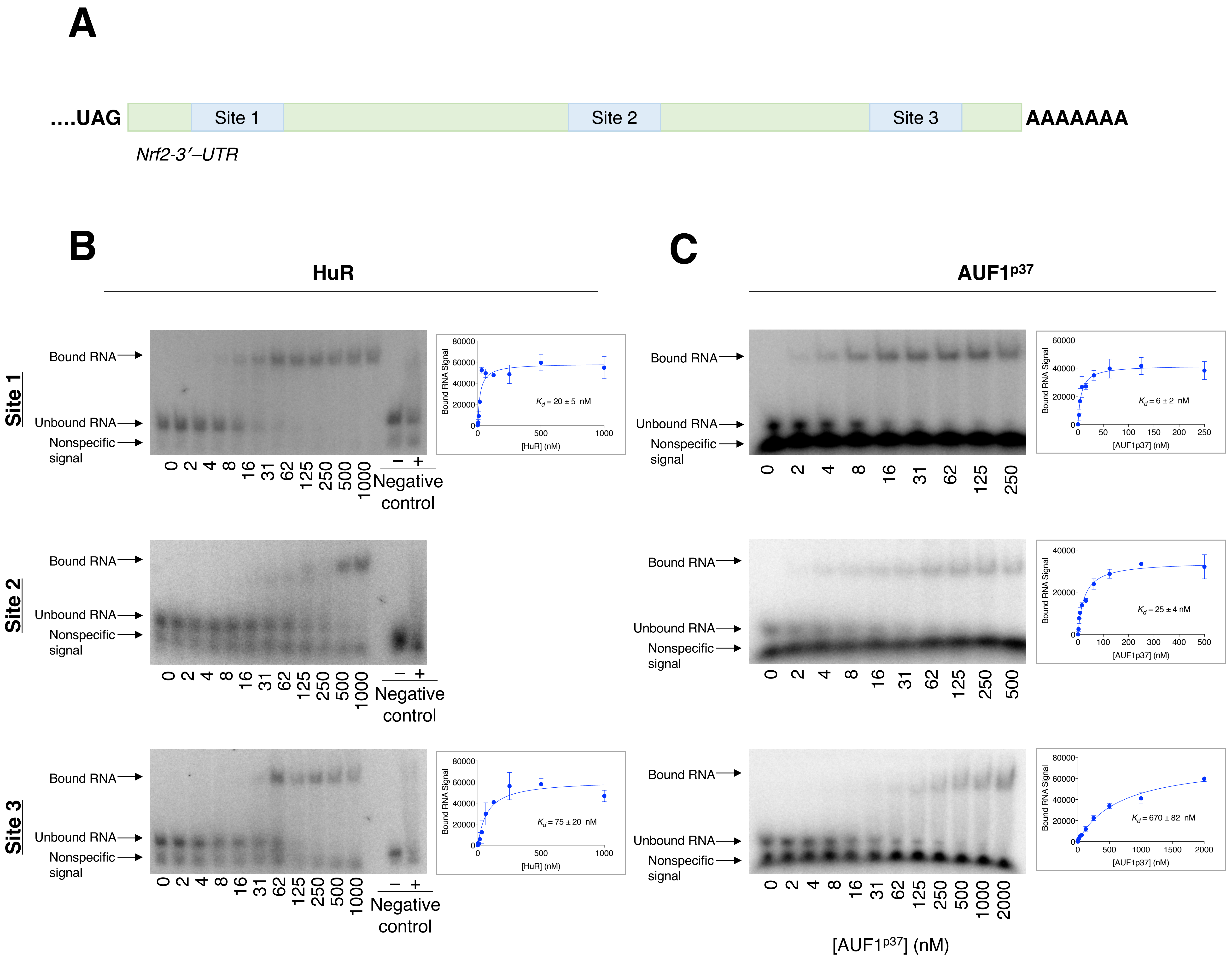

A
....UAG
Site 1
Site 2
Site 3
AAAAAAA
Nrf2-3′–UTR
B
HuR
C
AUF1p37
Site 1
Site 2
Site 3
Bound RNA
Unbound RNA
Nonspecific signal
0 2 4 8 16 31 62 125 250 500 1000
− +
Negative control
0 2 4 8 16 31 62 125 250
0 2 4 8 16 31 62 125 250 500
0 2 4 8 16 31 62 125 250 500 1000 2000
Bound RNA Signal
[HuR] (nM)
[AUF1p37] (nM)
Kd = 20 ± 5 nM
Kd = 75 ± 20 nM
Kd = 6 ± 2 nM
Kd = 25 ± 4 nM
Kd = 670 ± 82 nM
[AUF1p37] (nM)

### Figure S10

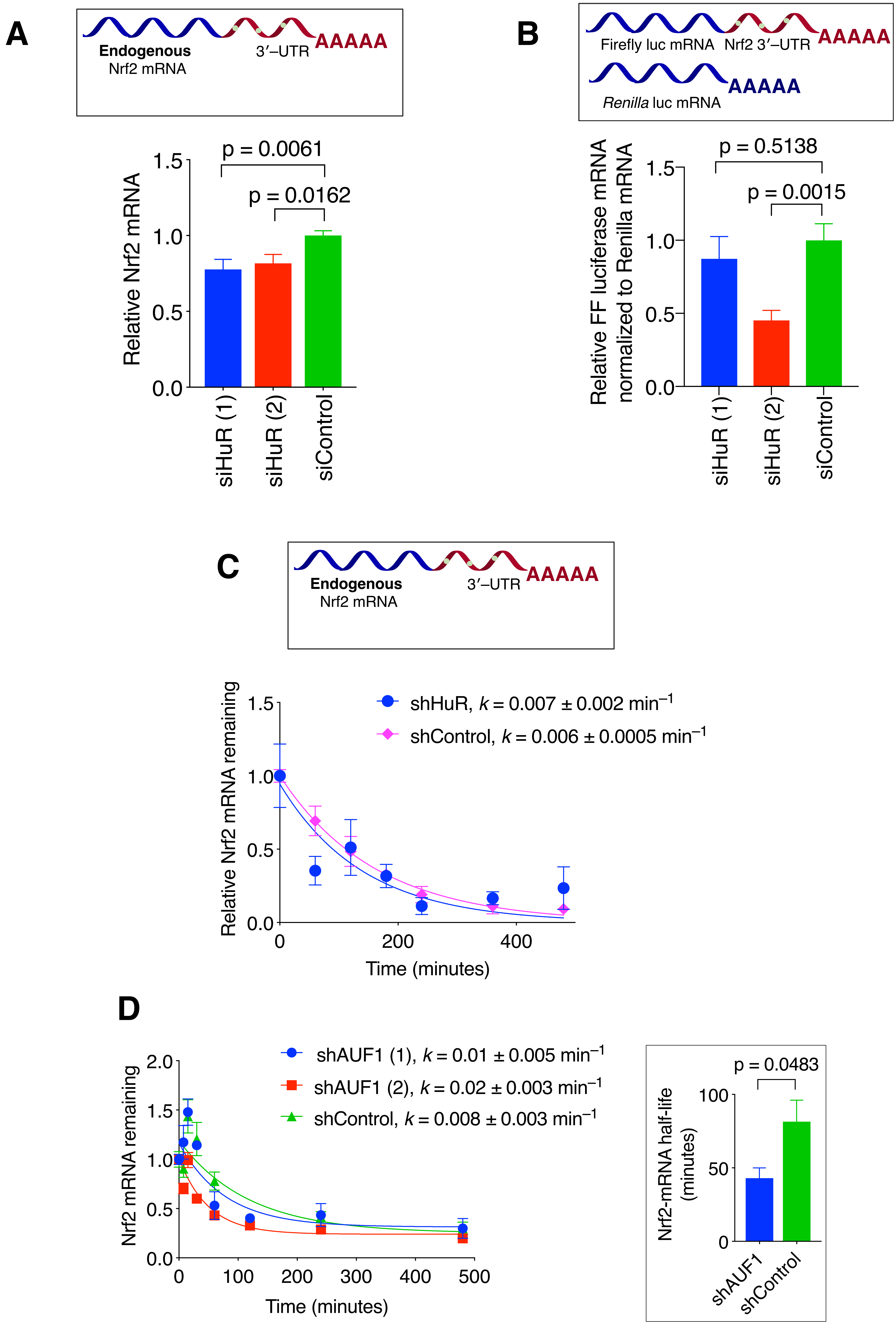

A
Endogenous
Nrf2 mRNA
3′–UTR
AAAAA
Relative Nrf2 mRNA
p = 0.0061
p = 0.0162
1.5
1.0
0.5
0.0
siHuR (1)
siHuR (2)
siControl
B
Firefly luc mRNA
Nrf2 3′–UTR
AAAAA
Renilla luc mRNA
AAAAA
Relative FF luciferase mRNA normalized to Renilla mRNA
p = 0.5138
p = 0.0015
1.5
1.0
0.5
0.0
siHuR (1)
siHuR (2)
siControl
C
Endogenous
Nrf2 mRNA
3′–UTR
AAAAA
Relative Nrf2 mRNA remaining
shHuR, k = 0.007 ± 0.002 min–1
shControl, k = 0.006 ± 0.0005 min–1
1.5
1.0
0.5
0.0
0
200
400
Time (minutes)
D
Nrf2 mRNA remaining
shAUF1 (1), k = 0.01 ± 0.005 min–1
shAUF1 (2), k = 0.02 ± 0.003 min–1
shControl, k = 0.008 ± 0.003 min–1
2.0
1.5
1.0
0.5
0.0
0
100
200
300
400
500
Time (minutes)
Nrf2-mRNA half-life (minutes)
p = 0.0483
100
50
0
shAUF1
shControl

### Figure S11

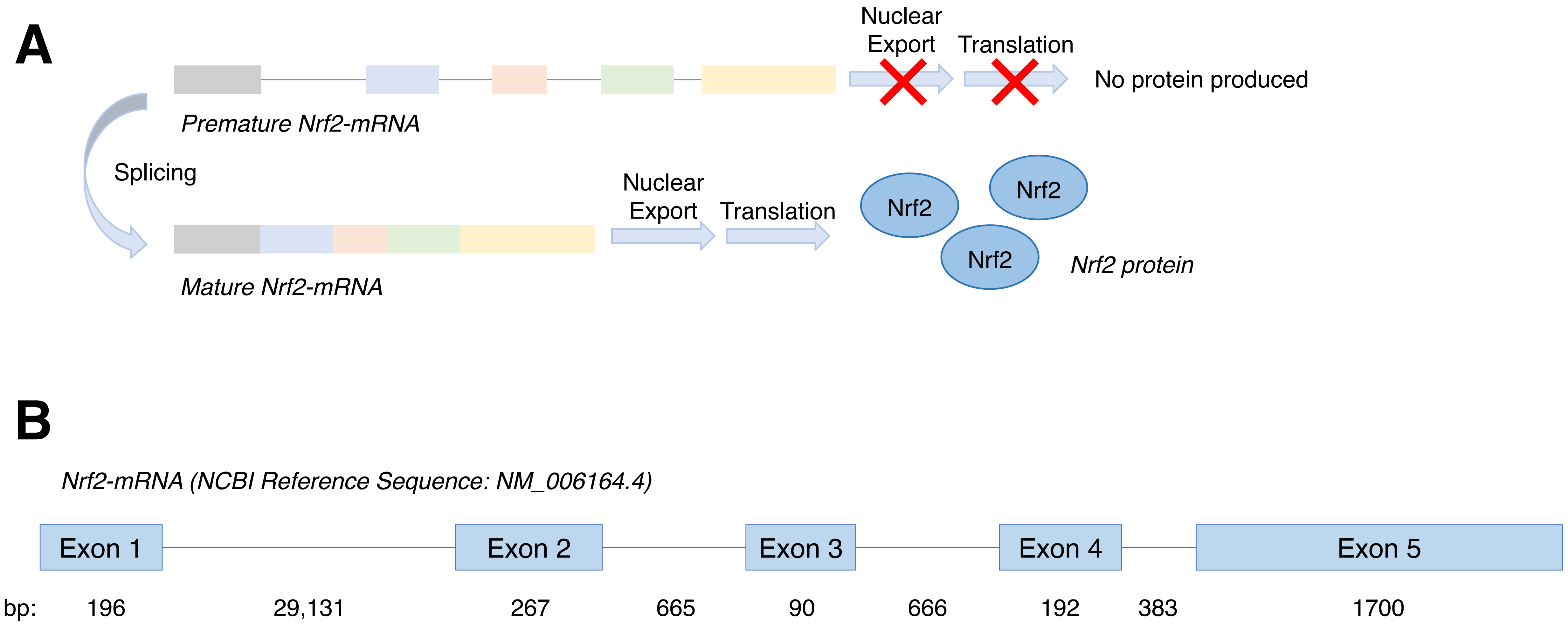
A
Nuclear Export
Translation
No protein produced
Premature Nrf2-mRNA
Splicing
Nuclear Export
Translation
Nrf2
Nrf2
Nrf2
Mature Nrf2-mRNA
Nrf2 protein
B
Nrf2-mRNA (NCBI Reference Sequence: NM_006164.4)
Exon 1
Exon 2
Exon 3
Exon 4
Exon 5
bp: 196 29,131 267 665 90 666 192 383 1700
